# Predicting rapid adaptation in time from adaptation in space: a 30-year field experiment in marine snails

**DOI:** 10.1101/2023.09.27.559715

**Authors:** Diego Garcia Castillo, Nick Barton, Rui Faria, Jenny Larsson, Sean Stankowski, Roger Butlin, Kerstin Johannesson, Anja Marie Westram

**Author notes:** Corresponding authors: Diego Garcia,; Anja Marie Westram. These authors contributed equally to this work (Roger Butlin, Kerstin Johannesson, Anja Marie Westram).

## Abstract

Predicting the outcomes of adaptation is a major goal of evolutionary biology. When temporal changes in the environment mirror spatial gradients, it opens up the potential for predicting the course of adaptive evolution over time based on patterns of spatial genetic and phenotypic variation. We assessed this approach in a 30-year transplant experiment in the marine snail *Littorina saxatilis*. In 1992, snails were transplanted from a predation-dominated environment to one dominated by wave action. Based on spatial patterns, we predicted transitions in shell size and morphology, allele frequencies at positions throughout the genome, and chromosomal rearrangement frequencies. Observed changes closely agreed with predictions. Hence, transformation can be both dramatic and rapid, and predicted from knowledge of the phenotypic and genetic variation among populations.

## Main text

Populations can sometimes adapt rapidly to sudden environmental shifts, even within a few dozen generations (*1, 2*). For many populations, rapid adaptation would be necessary to persist in the face of anthropogenic environmental change (e.g. climate change, habitat fragmentation, pollution, etc.). However, we are far from being able to predict whether and how fast a population will adapt, and which phenotypic and genetic changes will occur (*3*). We urgently need to understand whether adaptation is possible and how rapid it can be (*4*).

Adaptation relies on genetic variation, including both variation at individual base positions (*5*) and larger structural variants. The later include chromosomal inversions that generate large gene blocks that are inherited together and can simultaneously affect multiple traits (*6, 7*). Rapid adaptation particularly depends on variation already present within a species, because time is not sufficient to accumulate new beneficial mutations, unless population sizes are very large (*8, 9*) or generation times are very short (*10*).

The reliance of rapid adaptation on pre-existing variation suggests that it might be possible to predict future evolutionary change from knowledge about current variation (*11*). In particular, many temporal environmental changes, such as temperature increase, resemble a current pattern in space (e.g. a spatial temperature gradient). In this case, for a focal population experiencing an environmental change, adaptive evolution is likely to rely on genetic variation that has entered the population via past or on-going gene flow from a population that has already adapted to a similar environment. This information is often available from studies on phenotype-environment and genotype-environment associations. Can this knowledge on adaptive variation in space be used to predict how a population will respond over time after an environmental change? This principle is implicit in much conservation genetics work (*12*–*14*), but has rarely been explicitly tested. From a practical viewpoint, predictability would mean that population responses to environmental change can be anticipated and management efforts adjusted accordingly (*15*). In basic research, predictability provides a test of the current understanding of a system: for example, if loci contributing to divergence between environments in space have been identified correctly, they should respond in a predictable way to changing selection pressures in time.

The marine snail *Littorina saxatilis* is a model system in which divergent adaptation in space is exceptionally well-documented (*16*–*18*). Spatial variation and local adaptation to rocky shore environments are particularly obvious in the “Wave” and “Crab” ecotypes that have been intensively studied in Sweden, UK and Spain. The ecotypes originated repeatedly in different locations (*17*), in response to the selective pressures of wave action (*19*) and crab predation (*20*) on wave-exposed rocks and crab-rich parts of shores, respectively (*16, 21*) (Figure 1A). Adaptive variation in space in this system has been studied on three levels. At the phenotypic level, the ecotypes differ in traits like size, shell shape, shell colour, and behaviour (*16, 21, 22*). For example, the Wave ecotype is small, has a thin shell that often shows Wave-specific colours and patterns, a large and rounded aperture, and bold behaviour, while the Crab ecotype is large, has a thick shell without patterns, a relatively smaller and more elongated aperture, and wary behaviour (Figure 1B, Figure S1D). At the level of individual SNPs (single-nucleotide polymorphisms), highly differentiated loci likely contributing to adaptation or linked to adaptive loci are scattered across the whole genome (*18, 23*). At the level of large chromosomal rearrangements, several inversions differ in frequency between ecotypes (*24*–*26*) and explain variation in divergent traits between ecotypes (*23, 26*). These features all change over local contact zones between ecotypes and most differences are paralleled over large geographic areas (*27*). These repeated phenotype-environment and genotype-environment associations strongly suggest a role of divergent selection. Hence, we tested whether the observed spatial associations allow us to predict changes in time by studying evolution after an immediate environmental change in real time.

**Figure 1.**
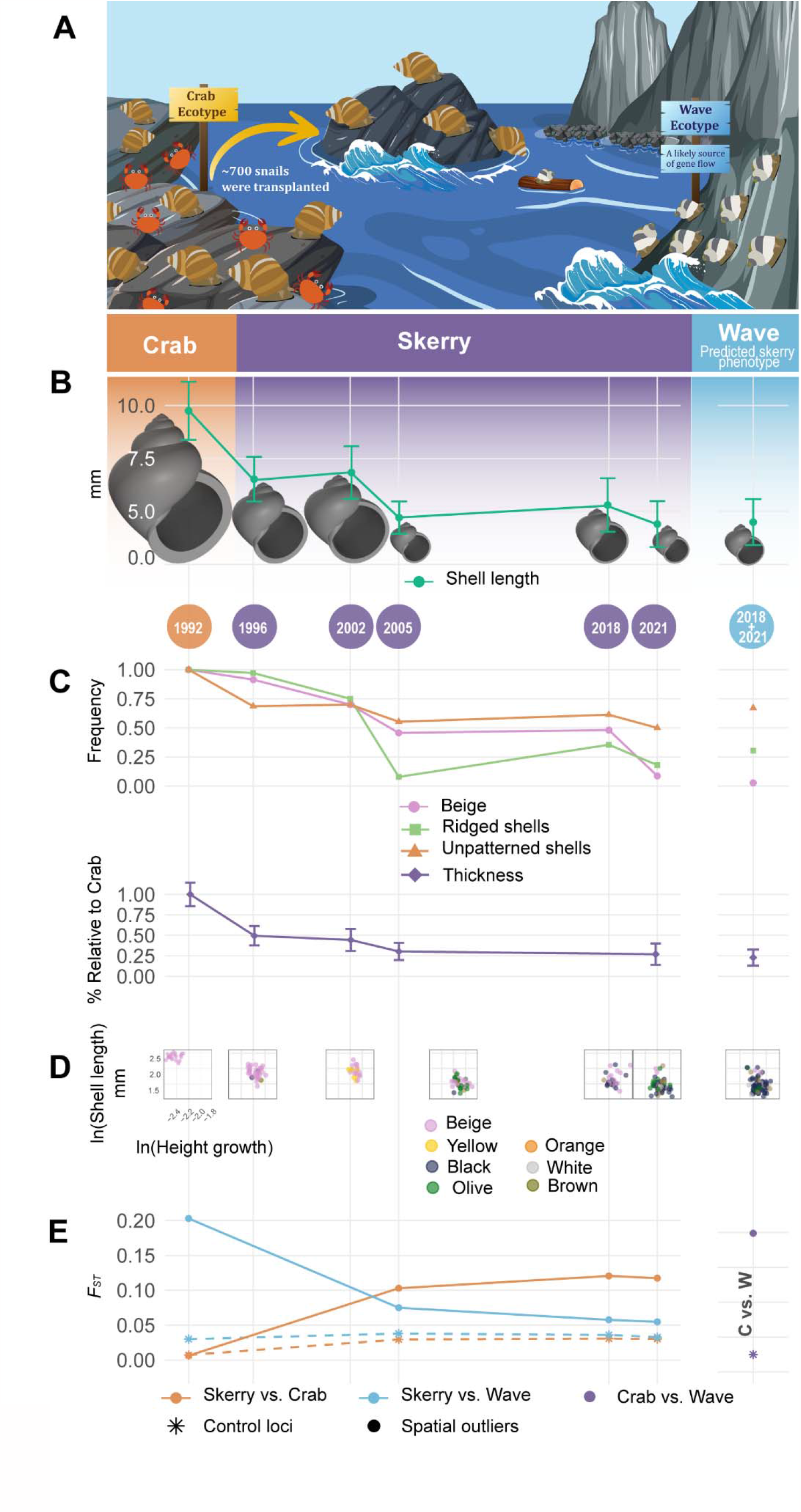
Divergence trajectory of the skerry population at the levels of phenotypes and loci in collinear genomic regions. Years correspond to sampling points in time. **(A)** Illustration of the transplant experiment showing the donor Crab ecotype on the left side, the recipient skerry in the middle, and the neighbouring Wave ecotype on the right side. Figure created using some graphics from Vecteezy.com under Free License. **(B)** Shell length and shape divergence. The shell length decreased after transplantation to less than half of the original value and approached the length of the reference Wave ecotype population. **(C)** Divergence of shell colour, patterning, roughness and thickness in the Skerry population that over time reached values in the range of the reference Wave ecotype. Thickness represent the average thickness in the Skerry population and Wave population relative to the average thickness of the transplanted population in 1992. **(D)** Scatter plots of two uncorrelated quantitative traits (shell length & height growth) on a log scale, and one qualitative trait (colour) reveal no bi-modalities in the skerry population. **(E)** Genetic differentiation of the skerry versus the reference populations based on control and spatial outlier SNP (single-nucleotide polymorphism) loci.

We assessed local adaptation in a 30-year transplant experiment on the Swedish west coast. In 1992, we collected ∼700 Crab ecotype snails and relocated them to a nearby wave-exposed environment earlier occupied by a population of the Wave ecotype. This wave environment is a “skerry” (a 1 × 3 m rocky islet), exposed to strong waves and with no evidence of crabs (Figure 1A; Figure S1). The skerry had remained uninhabited by snails since a toxic algal bloom in 1988 killed all Wave snails (*28*). The skerry (current census size: ∼1000 individuals) is located ∼320 m away, across open sea, from the donor Crab ecotype population and ∼160 m from the nearest Wave ecotype population (Figure S1; supplementary materials and methods). Therefore, there are two potential sources of adaptive variation: Standing genetic variation in the donor population (resulting, in part, from past gene flow from adjacent Wave populations on the same island), and post-transplant gene flow due to occasional migrants (e.g. rafted snails, see (*28*); the species lacks pelagic dispersal) from the neighbouring Wave population (or, less likely, elsewhere).

### Three predicted levels of adaptive evolution

We predicted three levels of change in the skerry population. At the phenotypic level, we anticipated a transition from Crab ecotype to Wave ecotype morphology: the averages of quantitative traits (e.g., shell length, and shell thickness) and the proportions of qualitative traits (e.g., shell colour, patterning, and roughness) were expected to approach the values typically observed in the Wave ecotype present in the area. We formulated our prediction on the basis of the polygenicity of phenotypes (*23, 29*) that can reach Wave optima through different pathways (both genetic and plastic) and are often under strong selection in space (*30*). For SNPs, we predicted an allele frequency shift over time beyond the effect of drift and neutral gene flow in at least a subset of “spatial outliers” (candidate SNPs associated with ecotype divergence in space in previous studies; see supplementary materials and methods) towards the frequencies observed in undisturbed Wave ecotype population. For inversions, we predicted an increase in frequency of arrangements that are common in Wave ecotype populations. We predicted a tendency to fix arrangements that appear favoured in the Wave ecotype (*18, 23*). We predicted non-fixation for inversions that are maintained polymorphic in the Wave ecotype, likely by balancing selection (*24*). Finally, for both spatial outlier SNPs and inversions, we predicted a correlation between temporal (start vs end of experiment) and spatial (Crab ecotype vs Wave ecotype) genetic differentiation. Overall, we expected predictability to be higher for inversions than for SNPs because many inversions are likely to be under strong direct selection, while spatial outlier SNPs may often only be indirectly affected by selection.

### Swift transformation of shell morphology and patterning validates phenotypic predictions

To evaluate our predictions at the phenotypic level, we sampled adult snails from the skerry in 1996, 2002, 2005, 2018 and 2021. As anticipated, the morphology of the transplanted snails experienced multiple changes from its original Crab ecotype to a morphology more similar to the Wave ecotype. In addition to a decrease in length, a shell reconstruction using six shape parameters (Figure 1B; supplementary materials and methods) revealed that, after 30 years, snails of the skerry population exhibited a relatively broader aperture and less pointed tips compared to their ancestral form and similar to the Wave ecotype. Moreover, the beige colour common in the Crab ecotype became rare over time, with the skerry population becoming colour-polymorphic, similar to Wave ecotype populations (Figure 1C). Simultaneously, the distinctively thick, ridged, and unpatterned shells of the Crab ecotype were largely replaced by thinner, smoother, and tessellated shells. Scatter plots depicting diagnostic traits (Figure 1D) show that this transition took place across all sampled individuals. Therefore, the change in population average on the skerry did not reflect the presence of migrants or early-generation hybrids but was due to gradual allele frequency changes across the whole population.

Previous estimates of additive genetic variance and plastic effects of the environment for size and shell-shape traits allowed us to estimate the strength of selection required to explain the observed phenotypic (*23*) changes in these quantitative traits. Assuming an initial plastic response in the transplanted population, followed by Gaussian stabilizing selection towards an optimum defined by the phenotype of the Wave reference population (supplementary materials and methods), the strength of stabilizing selection (V_S_/V_P_, the variance of the fitness function relative to the phenotypic variance) ranged from 1.65 to 7.84, depending on the trait, assuming one generation per year (supplementary text). These values are in the typical range for estimates from natural populations (*31, 32*) and correspond to a fitness reduction for the Crab ecotype population, when first introduced to the skerry between 30 and 96% (8 to 90% after the plastic change in phenotype). For the aperture position trait (r0) all of the change on the skerry could be accounted for by plasticity; for other traits, estimated plastic effects accounted for 23-40% of the change in phenotype on the skerry (Table S2).

Multi-year genetic data confirm adaptive frequency shifts in candidate mutations To evaluate our predictions at the genetic level, we genotyped samples from different years (2005, 2018, and 2021) from the skerry population as well as from the donor Crab population (1992, 2018, and 2021) and the neighbouring Wave ecotype population (2018 and 2021) (supplementary materials and methods). We included both spatial outliers (292 SNPs) that showed high Crab-Wave differentiation in previous studies on ecotype differentiation in the area (*18, 27*), SNPs diagnostic for chromosomal rearrangements (225 SNPs) (*25*), and control SNPs (565 SNPs) that lacked strong association with ecotype divergence in Sweden (*18, 27*). All spatial outliers and control loci are SNPs outside chromosomal inversions.

At the level of individual loci, the allele frequencies at many control loci in the skerry population changed towards the frequencies observed in the Wave ecotype: 59% of the control loci had shifted towards Wave in 2005, 63% in 2018, and 61% in 2021. For spatial outlier loci, the shift toward Wave was stronger, as predicted: 82%, 87%, and 89%, respectively (Figure S13, S14). The trajectory of genetic differentiation (*F*_*ST*_) also reflected the more pronounced shift of spatial outliers compared to the control loci (Figure 1E): The trajectory of spatial outliers indicate that from 2005 onwards, the skerry population was highly divergent from the Crab ecotype but close to the Wave ecotype. Control loci, on the other hand, showed no trend in direction in *F*_*ST*_. The results are also confirmed by a PCA (Figure S15).

The subtle change in allele frequency observed at control loci towards the Wave ecotype suggests either gene flow from a nearby Wave population or hitch-hiking effects by spatial outliers under selection. The fact that the allele frequency shift is more pronounced at spatial outlier loci than control loci is consistent with selection, although it alone does not provide sufficient evidence. This is because spatial outlier loci are on average more differentiated than control loci between the skerry starting population and the nearby Wave population. Therefore, neutral gene flow from a Wave population alone would already lead to a more pronounced shift for spatial outlier loci over time on the skerry population. To distinguish between these possibilities, we inferred the demographic history of the skerry population and compared the observed allele frequency changes to those expected under neutrality (neglecting linkage disequilibrium). Based on the allele frequencies of the control SNPs in the starting population (1992) and in the neighboring Wave population (2018+2021) as a potential source of gene flow, we found the growth rate, carrying capacity, migration rate, and number of generations per year that best predicted the allele frequency distribution observed in the skerry samples from 2005, 2018, and 2021 (supplementary materials and methods). The most likely estimate for migration was M=2 diploid individuals per generation (see Table S7 for other parameters). This relatively low number of migrants is reasonable considering that the species is brooding and without pelagic larvae and that the skerry remained unoccupied by snails for four years after the toxic algal bloom (*28*). Furthermore, it is of the same order of magnitude as direct estimates of colonisation of empty skerries in the area following the bloom (*28*). Starting with the allele frequencies observed in 1992, and randomly drawing parameter combinations from the likelihood surface, we simulated neutral evolution for each control and spatial outlier locus until 2021 (approximately 58 generations). Running 1,000 replicates for each SNP, we estimated the expected range of allele frequency changes from 1992 to 2021 without selection, but including genetic drift, gene flow, sampling, and model uncertainty (range spanning 95% of the simulated replicates) (Figure S12). While for both control loci and candidate outliers the probability of being outside the expected range is relatively small (4 vs 8.5%), the candidate outliers overall clearly show stronger allele frequency changes than expected under the neutral model (supplementary text). In alignment with our predictions, spatial outliers shifted towards Wave more than expected without selection (71% showed a stronger shift than the median shift without selection, compared to 53% in the control loci) (Figure 2A). This supports our prediction that selection influences at least a subset of the spatial outliers. Given that selection is not expected to directly impact control loci, drift, gene flow, and hitch-hiking effects are plausible reasons for the towards-wave shift in allele frequencies experienced by about half of the control loci.

**Figure 2.**
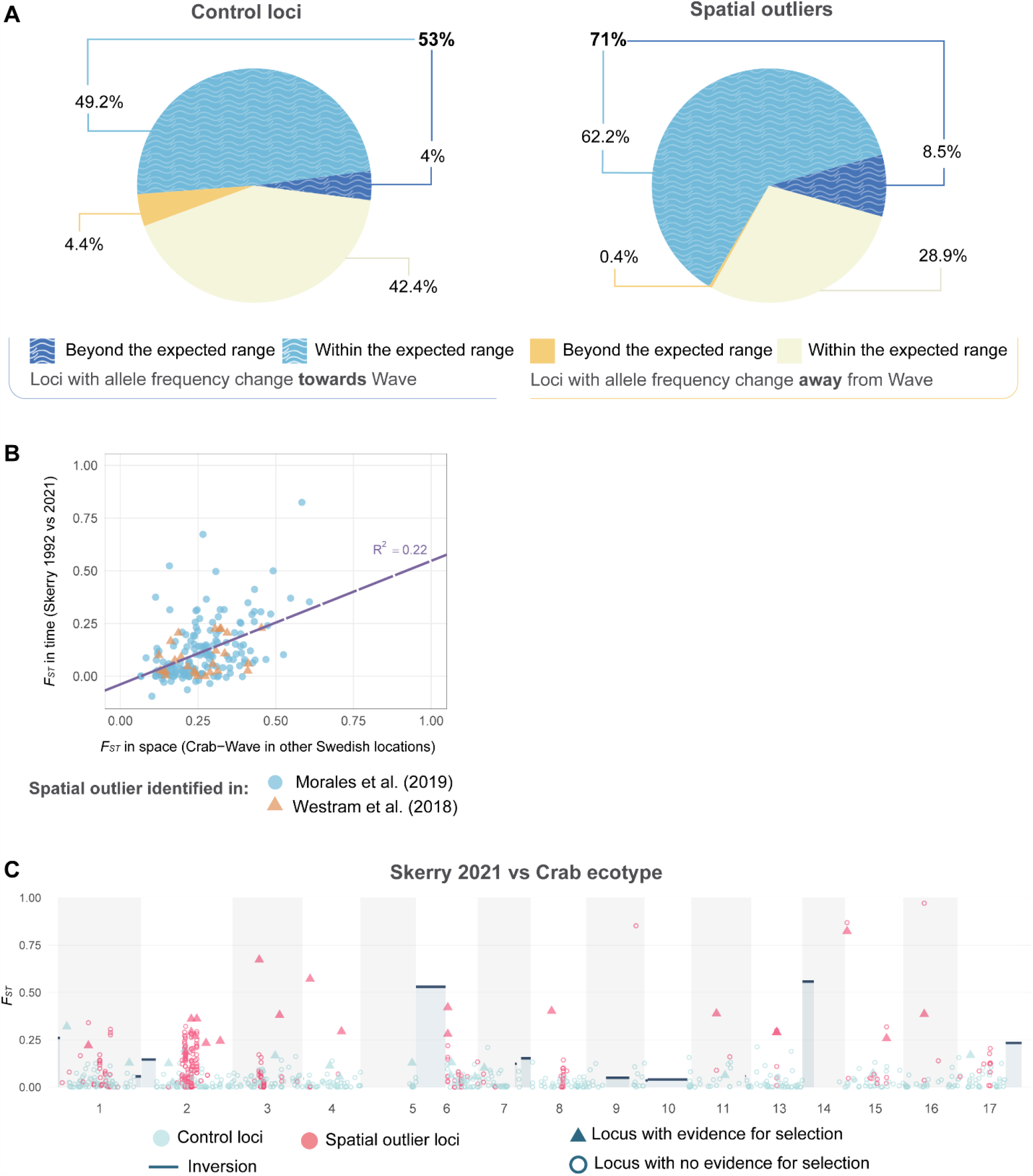
Genome wide evidence of selection in the skerry. **(A)** A comparison to the expected 95% range of allele frequency changes without selection shows evidence of selection in a larger fraction of spatial outliers than in control loci (dark blue slices of the pie charts). **(B)** Relationship between spatial and temporal differentiation. Each point is a spatial outlier SNP identified in a previous study, and the spatial *F*_*ST*_ reflects the average across (Morales et al. (2019) outliers; light blue) or 7 (Westram et al. (2018) outliers; orange) nearby Crab-Wave contacts (supplementary materials and methods). The line reflects a linear model describing the relationship (R^2^=0.22). **(C)** Genomic distribution of *F*_*ST*_ in the skerry population versus the average Crab ecotype for individual SNPs in the collinear genome (circles and triangles) and for inversions (rectangular blue-grey fields with dark blue bars at top indicating *F*_*ST*_ value). Loci with evidence for selection (triangles) are SNPs that experienced allele frequency change in the Skerry population towards Wave and beyond the neutral range.

As predicted, the more differentiated a spatial outlier was between Crab and Wave in other locations in Sweden, the more its allele frequency also changed in time; however the relationship was noisy (Spearman’s rho = 0.46, p<0.0001; Figure 2B). Loci potentially affected by selection are distributed widely along the genome rather than concentrated in a few linkage groups (LG; Figure 2C; Figure S16). We observed relatively few spatial outliers with evidence for selection (red triangles) in LG 2 compared to the expectation from its density of spatial outliers (Figure S17).

### Natural selection favours specific chromosomal rearrangements as predicted

Consistent with our predictions, the “Wave arrangements” (inversion arrangements that are more common in the Wave ecotype than in the Crab ecotype) increased in frequency in the skerry over time to near fixation or to a frequency similar to that in the Wave population (Figure 3A). This pattern was found in “simple” inversions (where two alternative arrangements exist) and complex inversions (where three different arrangements exist; Figure 3B). The two complex inversions are known to be particularly strongly associated with Crab and Wave ecotype divergence and to influence adaptive traits (*18, 23*). Simulations of neutral expectations (Figure S11) show that arrangement frequency shifts required selection in five cases (simple inversions, LGC1.1, LGC10.2 & LGC17.1, and both complex inversions, LGC6.1/2 & LGC14.1/2). Moreover, the inversions showed growing genetic differentiation *F*_*ST*_ over time in the skerry population with respect to Crab, comparable to the differentiation observed in spatial outliers (Figure 2C, Figure S16).

**Figure 3.**
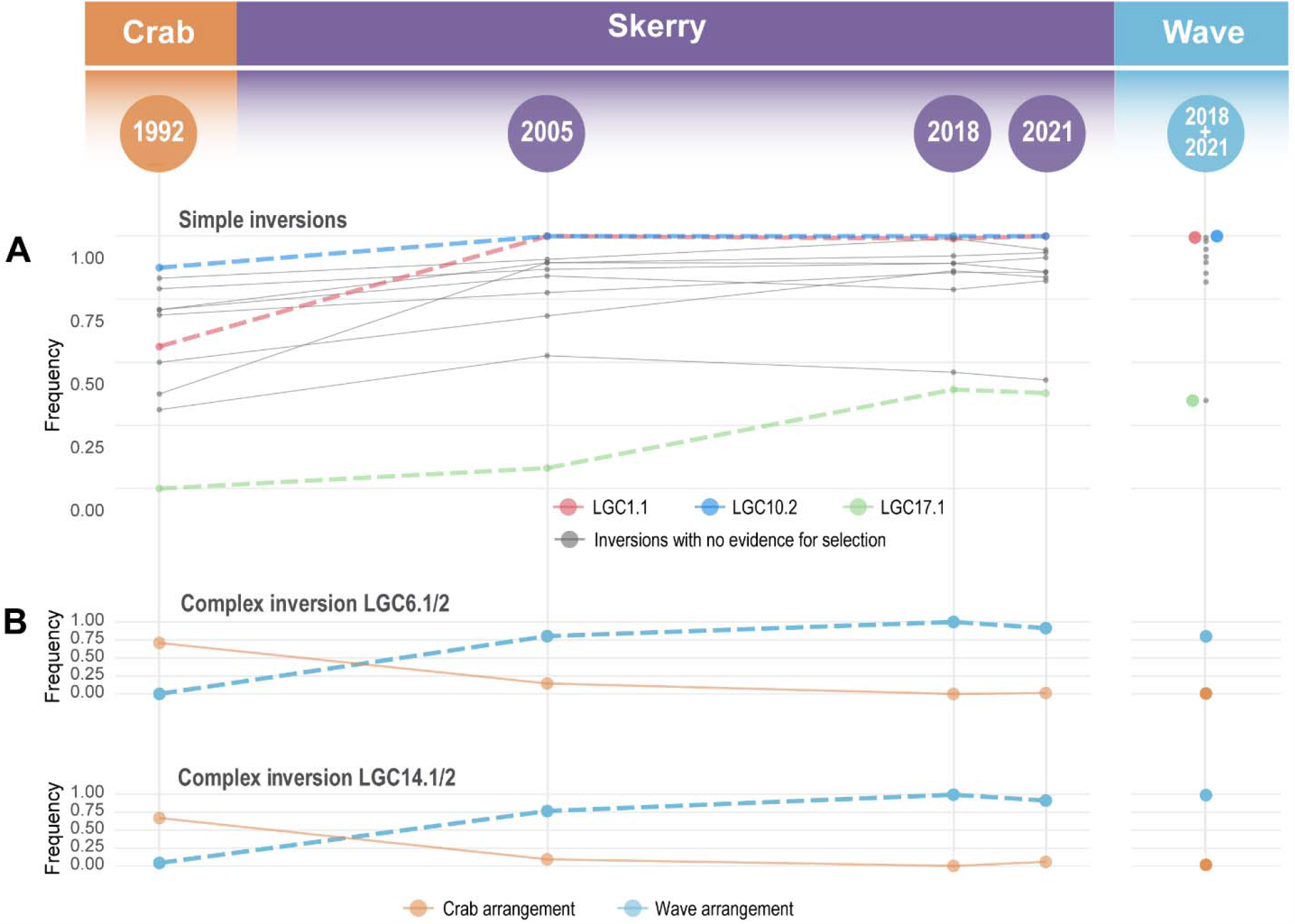
Trajectories of the “Wave” arrangement for polymorphic inversions. Grey lines indicate trajectories of arrangements in the skerry with frequency changes between 1992 and 2021 within the range expected without selection. The trajectories of arrangements that experienced frequency changes beyond the expected range are shown in colour. **(A)** The trajectories of simple inversions show arrangements of three inversions (LGC1.1, LGC10.2, & LGC17.1) that changed in frequency more than expected by drift and gene flow alone. **(B)** The “Wave” arrangement in complex inversions (dashed line) on two linkage groups (LG 6 AND LG 14) increased from rare to near fixation while the “Crab” arrangement (solid line) became rare.

### Discussion

This study shows that rapid phenotypic and genetic changes in a new environment can to a large extent be predicted based on spatial patterns. In barely a decade after the introduction, a transplanted population of Crab ecotype adapted rapidly to its new environment which closely resembles that of the Wave ecotype. Adaptive capacity following a sudden environmental change has been studied in other systems, such as guppy (*33*), stickleback (*2*), and salmonids (*34*). As in those studies, we demonstrated that adaptive evolution can take place relatively quickly, and we show that this is the case at three different levels: – phenotype, SNP genotype, and chromosomal inversion. Additionally, a morphological change immediately following the transplantation suggests that plasticity was crucial to avoid immediate extinction, as the allele frequencies adjusted to a new optimum (*35*).

Our results suggest that it is possible to predict how a population may change over time, using prior information on the genetic and phenotypic variation along spatial environmental gradients. However, predictability is only high at the phenotypic level and for strongly-selected inversions. Phenotypically, the skerry reached a Wave ecotype endpoint through the contribution of both adapted alleles (regardless of their source) and plastic changes. At the inversion level, the Wave arrangement in all cases reached frequencies similar to that in a Wave ecotype population, with statistical evidence for selection in five cases. This is in line with their widely accepted role in suppressing recombination between beneficial alleles in a specific genetic background or environment. Predictability is lower for collinear loci, for which only a small fraction showed clear evidence of selection and this specific subset could not be predicted. There are multiple, not mutually exclusive explanations. First, some loci might be less favoured on the skerry than in the wave-lashed environment that we used as a reference. Thus, these loci might be under selection only in some wave habitats because the environmental conditions differ among locations (e.g., the Wave samples in previous studies are likely to be associated with higher shore levels than is the skerry). Second, many of our spatial outliers may be linked to targets of selection, rather than being under selection themselves; in this case, responses to selection depend on the linkage disequilibrium in the studied population, which in turn depends on the history of gene flow. Third, because some of the adaptive traits are likely to be highly polygenic (e.g. shell shape), the same phenotypic optimum can potentially be reached via different genetic routes (*29*). Therefore, it is plausible that the evolution of the Wave phenotype was possible via changes at a subset of the loci studied here, together with changes at loci not included in this study.

A recurring challenge for genomic studies of this nature consists of disentangling the effects of demographic history and natural selection (*36*). For example, if alleles introduced by migration experience positive selection and large blocks of migrant (Wave) background hitchhike along, we might overestimate the migration parameter. However, this does not affect our result that spatial outliers and inversions shift more towards Wave than expected without selection. In addition, we did not find evidence that hitch-hiking extends over a larger region of the chromosomes (Figure S18).

In the Anthropocene, studies such as the present one can be a basis for developing predictive models for the response of populations to environmental changes as a result of human activities, e.g. climate change, industrial fishing, habitat fragmentation, invasions, etc. (*37*). Future experiments could integrate variables (e.g. temperature, precipitation, pollution, etc.) that fluctuate as a result of human activities. In our study, we observed rapid adaptation, and predicted the genetic and phenotypic changes successfully because the population experiencing an environmental change contained or received a great amount of genetic variation, the raw material for natural selection. Nonetheless, this scenario will not universally apply to numerous other populations undergoing (anthropogenic) environmental shifts. Our results highlight the importance of ensuring that species remain in a variety of different environments in order to maintain genetic variation needed to fuel future adaptation.

## Supporting information

Supplementary Table S2 Phenotypic Analysis

Skerry interpolation 9.23 v2

## Author contributions

Conceptualization: AMW, KJ, RKB

Analysis: AMW, DGC, JL, NB, RKB, SS

Writing – original draft: DGC

Writing – review & editing: AMW, DGC, JL, KJ, NB, RF, RKB, SS

## Data and materials availability

SNP and phenotype datasets will be available on a repository upon acceptance.

