## Supplementary Material for "Predicting rapid adaptation in time from adaptation in space: a 30-year field experiment in marine snails"

**Affiliations:**
[1]Institute of Science and Technology Austria (ISTA), Klosterneuburg, Austria
[2]CIBIO-InBIO, Centro de Investigação em Biodiversidade e Recursos Genéticos, InBIO, Laboratório Associado, Universidade do Porto, Vairão, Portugal
[3]BIOPOLIS Program in Genomics, Biodiversity and Land Planning, CIBIO, Campus de Vairão, 4485-661 Vairão, Portugal
[4]Department of Marine Sciences - Tjärnö, University of Gothenburg, 45296 Strömstad, Sweden
[5]Ecology and Evolutionary Biology, School of Biosciences, The University of Sheffield, Sheffield S10 2TN, UK
[6]Faculty of Biosciences and Aquaculture, Nord University, 8049 Bodø, Norway
* Corresponding authors.
† These authors contributed equally to this work

**This PDF file includes:**

Materials and Methods
Supplementary Text
Figs. S1 to S18
Tables S1 to S9

**Other Supplementary Materials for this manuscript include the following:**

Table S2: Supplementary Table Phenotypic Analysis.xlsx
Mathematica Workbook: Skerry interpolation 9.23 v2.nb

### Materials and Methods

Description of the sites, the translocation and the subsequent sampling

The study area is part of the Kosterhavet National Park on the Swedish west coast. It is an archipelago of hundreds of small islands and rocks (Figure S1). The skerry is a tiny isolated rock: 1 x 3 m wide, and 0.5 m height over mean water level (58.84128N, 11.05111E). It is separated by 160 m of 5-20 m deep water from the nearest island. The area has only a very restricted tidal range (max 0.35 m) and the rock is only occasionally completely submerged. In 1985 a population of about 1000 Wave ecotype *Littorina saxatilis* inhabited the skerry. In May 1988 the population was wiped out by a unique bloom of a toxic microalgae (Chrysochromulina polylepis) (Johannesson & Johannesson 1995) and visits in June and October the same year and in summer 1989 confirmed that the snail populations was completely extinct on the skerry.

A visit in May 1992 again showed that no recolonization had taken place, and we decided to make a translocation of Crab ecotype snails from a nearby donor population. This donor site (58.84397N, 11.05279E) is a nearby boulder shore 320 m away from the skerry. It is densely populated by *L. saxatilis* of Crab ecotype over 100 m of shore, which means roughly >20,000 adult snails. This Crab ecotype population is flanked by snail populations of the Wave ecotype on both sides. On 30 May 1992 approximately 600-800 adult Crab ecotype snails were moved from the donor site to the skerry. A subsample of Crab ecotype from the donor site was stored as a reference. One month later only 50 of the translocated snails could be observed on the skerry, but most likely approximately several hundred 0.5 mm juveniles (invisible to the naked eye) were already released on the skerry. (The species is brooding, and females are fertile year-round and release on average 1-2 juveniles per day).

Sampling of adult snails from the skerry was undertaken in 1996, 2002, 2005, 2018 and 2021. At all occasions only a minor proportion of the total population was sampled, typically about 30-35 snails out of 1000. New samples were also taken from the donor site in 2018 and 2021 and from a Wave ecotype reference population (58.84066N, 11.04794E) 160 m west of the skerry. The reference site represents the closest Wave ecotype population to the skerry, and represents one of several nearby wave populations from which migration of single snails by rafting could have taken place. Indeed, following the extinction of all snails on intertidal skerries in the area, after 4 years, 27% of the skerries had received migrants and in 12% of the skerries population sizes were restored (Johannesson & Johannesson 1995). While skerries only a few meters from islands had higher migration rates than more remote skerries (like the experimental skerry of the current study), this still indicates a potential for a small number of migrants to arrive by rafting from surrounding populations, in addition to the translocated snails from the donor population between 1992 and 2021.

Phenotyping

Phenotyping of shell traits included shell length, shell thickness (mean of three measurements per snail), shell colour and pattern, shell ornamentation and size-independent parameters for shell shape. These measurements were obtained from photos, measured as described in Larsson et al. (2020), and analysed using a growth model as described in such a study. The same measurements were done on all samples with the exception that shell thickness was not measured on the 2018 samples. Shells were categorized into seven colour groups: beige, yellow, black, olive, orange, white, and brown. The classification as either tessellated or ridged was determined by the presence or absence of ornamentations (such as stripes) and striations on the shell, respectively *[Phenotype-datasets will be available upon acceptance]*.

SNP development and genotyping

We developed a panel of SNPs and genotyped our samples using a targeted genotype-by-sequencing service provided by LGC Bioresearch Technologies. We first identified a large number of SNPs (20,000) based on previous genomic studies of Crab and Wave ecotypes in *Littorina saxatilis.* These included (i) 'control' SNPs that show no prior evidence for selection between the ecotypes (*27*), (ii) SNPs that show selection based on cline analysis across a single Crab-Wave transition (ANG) (*24*), (iii) SNPs that show evidence for selection in a Baypass outlier scan performed on multiple Crab-Wave contrasts across Europe (*27*), (iv), SNPs that show high differentiation (based on $F_{ST}$) in 1 or more out of 5 Crab-Wave contrasts in the Koster marine park in Sweden, and (v) SNPs that are diagnostic for chromosomal inversions that show evidence for a selection across a  Crab-Wave transition in Sweden (ANG)(*18, 24*, *25*).

From this large set of possible SNPs, we then selected 5000 for an initial genotyping trial. For the control SNPs, we chose 1 marker from each of 1000 map positions (acording to a previously published genome and genetic map (*24*)), and a minor allele frequency greater than 0.1. We chose 1161 Baypass outliers, biased toward markers with stronger evidence of selection (Most with Bayes factors of 2 or 3, with 3 being the strongest evidence). We also chose 16 diagnostic SNPs for each inversion, selecting those with the most power to distinguish the alternative arrangements. For the other categories, all potential SNPs were tested.

SNPs from this trial were carried forward if (i) they were successfully genotyped in at least 80% of individuals (with calls requiring at least 8 reads) and (ii) they were bi-allelic, single position variants. In the second trial, 24 individuals were genotyped using a different annealing temperature to improve probe hybridization. In addition to the above criteria, we only retained SNPs were the two alleles in heterozygotes where at a ratio between 0.3 and 0.7. The final selection of SNPs was chosen to maximise spread across the genome, including representation of collinear and inverted regions. We excluded all SNPs in LG 12, a region that has been associated with sex determination (23), and the majority of SNPs in LG 5, which might be part of a putative inversion (25).

As some of the categorization of SNPs into "control" and "spatial outlier" SNPs relied on Europe-wide data (*27*), we further filtered the collinear SNPs to obtain a set of control loci and spatial outliers that more accurately reflect the local patterns in Sweden. Using the poolseq data from Morales et al. (2019), we calculated a "local" $F_{ST}$ by averaging $F_{ST}$ across the four nearby Swedish locations (Arsklövet, Saltö, Ramsö, Jutholmen). If for a given location the same allele was fixed in Crab and Wave, $F_{ST}$ was set to 0 for that location before averaging. We then calculated the 95% quantile of the $F_{ST}$ distribution across loci. For the spatial outliers, we kept those that were above this quantile; for the control SNPs, we kept those that were below. The other SNPs were completely discarded. For the spatial outliers from Westram et al. (2021), we kept all, as the study was based entirely on nearby locations in Sweden.

All aspects of the DNA extractions and genotyping were performed by LGC Bioresearch Technologies. The final counts of SNPs retained across each category in this study are provided in Table S1 *[SNP datasets will be available upon acceptance]*.

Filtering

We used vcftools to filter the SNP dataset as follows; minimum SNP quality of 40 (--minQ 40), exclude indels (--remove-indels), exclude all genotypes with a quality below 20 (--minGQ 20), keep only genotypes with at least 10 reads (--minDP 10), keep only biallelic SNPs (--min-alleles 2 --max-alleles 2), and keep only sites with a minor allele count of 5 (--mac 5). Additional filters were applied in R. Before analysing collinear regions (neutral and outlier loci), we excluded SNPs with more than 5% missing data within one population and year, as well as individuals with more than 5% missing genotypes. For inversions, we excluded SNPs with more than 20% missing data within one population and year, as well as individuals with more than 20% missing genotypes. For PCA analysis of both collinear loci and inversions, we inputted the missing genotypes with the most common genotype within each population and year.

Shell reconstruction

The description of shell shape used in this paper is based on a logarithmic helicospiral growth model developed specifically to capture the shape variability present in the shells of different ecotypes of the snail *L. saxatilis* (Larsson 2020). Parameter values representing shape and growth are inferred from 2D images of shells in a standardised orientation and are used as quantitative morphological measurements. Since these parameters represent an approximation of the shell construction process, they give biologically relevant shape descriptions and enough information to generate 3D models of the shells.

In this analysis we used six parameters to quantify the shell shape (Figure S2), which have previously been found to correlate with the Crab-Wave ecotype differentiation in *L. saxatilis*. The parameter values were obtained using the program ShellShaper (https://github.com/jslarsson/ShellShaper), which infers the values from user input in the form of reference points and curves placed onto a standard orientation shell image. The program allows the user to visualise the 3D model generated by the obtained set of parameters together with the original shell, to directly compare their shapes.

Strength of selection based on phenotypes

The strength of selection acting on phenotypic traits was estimated for shell length and for five components of shell shape variation (gw, gh, a0, r0 and c; width growth, height growth, aperture radius, aperture position and aperture shape, respectively) based on the growth model of Larsson et al. (2020). Growth parameters a0 and r0 were expressed relative to shell length. Parameter gw was analysed without transformation, as in Koch et al. (2022), gh, r0 and shell length were log transformed.

For each phenotypic variable, we assumed stabilizing selection around an optimum phenotype, O:

$$w = e^{\frac{(x-O)^2}{2V_s}}$$

where $w$ is the fitness of an individual of phenotype $x$ and $V_s$ is the width of the fitness kernel. The expected change in mean phenotype in one generation is then:

$$\Delta x = (O - \bar{x})\frac{V_g}{(V_s + V_p)}$$

(Charlesworth & Charlesworth 2010, p.186), where $V_g$ is the additive genetic variance in the trait and $V_p$ is the phenotypic variance. Offspring of individuals introduced to the skerry would have had Crab genotypes but their phenotypes would have been influenced by plastic responses to the skerry environment. Therefore, our expectation for the phenotypic mean of the starting population on the skerry was $\bar{x}_{crab} + p$, where $p$ represents the plastic effect.

In order to fit this model to the available phenotypic data, we assumed that the optimum on the skerry is the same as the optimum phenotype in the environment of the Wave reference population and can, therefore, be estimated by the phenotypes of individuals sampled from that population. We used data from three nearby transects analysed by Koch et al. (2022) to provide estimates of $p$ and $V_g$. Specifically, environmental effect estimates by Koch et al (2022). were used to predict the change in phenotype for a Crab individual placed in the Wave habitat in the lower half of the shore height range. The mean of the estimates for the three sites and the variance among sites were used to provide a Gaussian prior distribution for $p$. Similarly, a prior for $Vg$ was set using the background additive genetic variance and inversion effects estimates from Koch et al. (2022). We used either estimates from the Crab populations only ($V_g$-*crab*) or from the whole transects ($V_g$-*site*) in separate analyses in order to bound the likely range of genetic variation available on the skerry, which derives from the Crab introduction plus input from migrant Wave individuals. We assumed either one or two snail generations per year: the average generation time probably lies between these two values. Estimates in the main text are for one generation per year and $V_g$-*crab*. Estimates from all combinations are provided in Table S2.

The model was then fitted using R-Stan version 2.21.7 (*38*). All samples from the Crab donor population were combined to estimate the starting mean phenotype and both samples from the Wave reference were combined to estimate the Wave optimum phenotype. Predicted phenotypes were fitted to the Skerry samples. The phenotypic variance, $V_p$, was assumed to be constant across all samples, with a Gaussian distribution. We checked this assumption by comparing samples using Levene's test and found no significant departure from homogeneity of variances with the scaling used. We fitted the ratio $V_s/V_p$ since this is comparable across phenotypes and we set a flat prior from 0 to 50 based on the values reported for natural populations (*31*, *32*). We also estimated the expected fitness of the mean Crab phenotype in the Wave environment. We used 4 chains of 3000 iterations, discarding the first 1000 iterations in each case as burn-in.

Analysis of phenotypes

The general pattern we observed in the phenotypic traits, both quantitative (such as shape parameters, length, and thickness) and qualitative (including ridging, colour, and tessellation), indicates that the Skerry population evolved a morphology similar to that of the Wave ecotype. However, we wondered how this transition from a Crab-like morphology to a Wave-like morphology occurred in the early stages. For instance, did this transition happen smoothly for the entire population, or was there a period during which more than one type of phenotype coexisted on the skerry, with some individuals exhibiting a more Crab-like appearance and others displaying a more Wave-like one?

We first identified the most informative quantitative traits: pairs of traits that are negatively correlated between the ecotypes (all data), but uncorrelated within an ecotype (Figure S3A). For example the pair width growth and average thickness. Second, among the candidate informative traits, we chose only those traits where there is relatively large difference between the Crab and Wave ecotypes. Our criterion was that we should not observe individuals of one ecotype within the interquartile range of the other ecotype (Figure S3B). Thus, the most informative quantitative traits are: thickness, shell length, height growth, and width growth.

We then plotted pairs of uncorrelated traits in a scatter plot colored by the three qualitative traits (Figure S4). We did not observe strong evidence for bimodalities in the scatter plots. Rather, the phenotypic transition of the skerry population occurred in the whole population. For a better visualization of the changing distribution of phenotypes over the years, we generated bar plots of the quantitative traits including qualitative data (Figure S5). The distribution of quantitative traits tend to shift towards Wave ecotype faster than qualitative traits. E.g. In 1996 the distribution of the shell length was shifted to the left but the frequency of ridged shells and the colour remained more or less the same as in the founder population.

<u>Demographic inference</u>

We inferred the demographic history of the skerry population using maximum likelihood. For that, we needed to calculate the probability of obtaining the observed allele frequencies in samples from the skerry, given a demographic parameters model. The set of model parameters that maximises this probability represents the most likely history of the population, and the set of parameters that have likelihood close to this maximum represents the plausible set.

Our main goals were: 1. To test whether we can exclude gene flow from the nearby Wave population, in which case adaptive change on the skerry must result from selection on standing genetic variation from the Crab donor. 2. To obtain the expectations for allele frequency changes under neutrality in order to test whether spatial outlier loci deviate significantly from this neutral model.

All analyses were run in R version 3.6.3, and for some parameters, an independent check combinations was run in Mathematica v. 12.3 (supplementary material Skerry interpolation 9.23 v2.nb).

*Data*

As we aimed to infer the neutral history of the skerry population, the demographic inference focused on control SNPs. To reduce effects of linkage, we only used the first control SNP from each contig (n = 438 SNPs). For the skerry, we had information from four sampling times: 1992 (when a sample from the Crab donor population was taken at the same time as collecting the donor individuals for the skerry), 2005, 2018 and 2021. The four allele frequency estimates on the skerry were the observed data whose probability we aimed to maximise. In addition, we needed information about allele frequencies in the Wave reference population to infer the extent of gene flow. As we assumed the Wave population to be stable over time (see Figure S15, PCA on collinear loci), we merged the 2018 and 2021 samples from the Wave population for this analysis.

To avoid missing data, we determined the minimum number of (diploid) individuals with data per SNP for each sample (Skerry 1992, 2005, 2018 or 2021; Wave 2018+2021) and then subsampled all other SNPs to that sample size. This led to the sample sizes shown in Table S3.

*Model*

We assumed a simple model of population growth with gene flow (Figure S6, Table S4). The model assumes that the skerry population was founded with $N_0$ haploid genomes (i.e. $\frac{N_0}{2}$ diploid individuals) sampled from the Crab donor population. We then assumed logistic population growth at rate $r$ with carrying capacity $K$ haploid genomes, where the population size at time $T$ is

$$N_T = \frac{K}{1+\left(\frac{K-N_0}{N_0}\right) exp(-rT)}.$$

There is unidirectional gene flow of $M$ haploid migrants per generation from the Wave population. We assumed non-overlapping generations, with $f$ generations per year, as the exact generation time of *L. saxatilis* for this population is unknown. Sampling therefore took place at the start of the experiment (1992), after $13f$ generations (2005), after another $13f$ generations (2018), and after an additional $3f$ generations (2021). Given that there cannot be fractions of generations in our model, each sampling generation was rounded to the nearest integer, and the total duration of the experiment was $round(13f) + round(13f) + round(3f)$ generations.

The parameter values tested are described in Table S4. We included a large range for $N_0$ because prior information about the starting effective population size is limited despite the known number of transferred individuals: on the one hand, it is likely that a considerable proportion of the transferred individuals did not attach to the rock and were immediately lost at the start of the experiment; on the other hand, because adult *L. saxatilis* females typically carry offspring from multiple fathers in their brood pouch, the effective starting population might have been larger than the number of adults. We also included a wide range of growth rates as little is known about the speed of population growth. The maximum carrying capacity $K$ included was based on the fact that ~2000 snails were estimated to live on the skerry in 2014, when the population was well-established. Numbers of migrants were capped at 8 per generation because the skerry remained empty for four years after the algal bloom, indicating that migration rates are unlikely to exceed a couple of migrants per year on average. The generation factor $f$ was based on generation times of *L. saxatilis* in the lab at similar temperatures (*39*).

*Inference*

To infer the most likely combination of parameters, we must calculate the probability of seeing the observed allele counts in the skerry sample for each combination of parameters.

The observed allele counts on the skerry at the different sampling times for a given SNP can be described by a vector $k = \{k_{1992}, k_{2005}, k_{2018}, k_{2021}\}$. For each sampling time, the probability of sampling $k_T$ depends on the real allele count at that time, $i_T$, in the population, and on the probability that the population has actually evolved to this count. The probability of sampling the vector $k$ is therefore

$$p(k) = \sum \psi_{92} s_{92} M_{92\to05} s_{05} M_{05\to18} s_{18} M_{18\to21} s_{21},$$

where the sum is over all possible combinations of $i_T$. This sum occurs because each $i_T$ is unknown and we therefore need to sum over all possible allele counts on the skerry.

$\psi_{92}$ is the prior distribution of allele counts in the starting population on the skerry. We assume a uniform prior, i.e. $\psi_x = \frac{1}{N_0+1}$ for each possible allele count $x$.

$s_T$ is the probability of sampling $k_T$ copies from the $i_T$ copies in the population at time $T$. This is a binomial sampling probability where the parameters are the sample size at sampling time $T$ and $i_T$.

$M_{T_a\to T_b}$ is the probability of evolving from $i_{T_a}$ to $i_{T_b}$ copies in the interval from $T_a$ to $T_b$. This is a transition matrix describing the evolutionary process on the skerry. This matrix is calculated as follows. Following basic population genetic principles, the expected allele count $i$ on the skerry after a single generation of migration from the reference Wave population is

$$i_{T_{exp}} = \frac{i_{T-1}}{N_{T-1}}(N_{T-1} - M) + p_{Wave}M.$$

From this, the probability of $i_T$ copies in the population at time $T$ can be calculated from a binomial distribution with parameters $N_T$ and $i_{T_{exp}}$ (the binomial sampling here represents the process of genetic drift). In the full matrix, the rows reflect all possible $i_{T-1}$, i.e. all possible allele counts in generation $T-1$ $(0,1,\dots N_{T-1})$, the columns reflect all possible allele counts in generation $T$ $(0,1,\dots N_T)$, and each cell gives the probability of evolving from $i_{T-1}$ to $i_T$ copies.

To obtain the negative log-likelihood for any parameter combination, we calculated $p(k)$ as described above for each SNP, took the negative log and summed across all SNP.

The best parameter combination from the parameter grid was defined as the one where the negative log-likelihood was minimised.

*Interpolation to find the truly best parameters*

Because only a relatively coarse grid of parameter combinations could be tested due to the computational demands of the analysis, we interpolated the likelihood surface to find the best parameter combination, including combinations not included in the grid. $f$ (the number of generations per year) was fixed at 2, as this value produced the highest likelihood, whilst the parameter overall seemed to have little effect (Figure S7; Figure S8; Figure S9; Table S5).

*Simulations to determine the "expected range" of allele frequency change without selection for each locus*

We ran simulations to test whether the allele frequency changes observed in the control SNPs and the spatial outlier SNPs are consistent with a neutral model, given our model uncertainity and sampling procedure.

For each SNP, we ran 1,000 replicate simulations, always using the allele frequency observed in Crab in 1992 and the allele frequency observed in the Wave reference for the respective SNP as input. For each simulation, the combination of demographic parameters was randomly sampled from the likelihood surface (described in the previous section). Each simulation started with the allele frequencies observed in the Crab sample in 1992 and included population growth (based on $r$ and $K$) and random binomial sampling under gene flow and drift (based on $M$, $p_W$ and the population size based on the growth parameters) in each generation. The simulations were stopped at the last sampling time point (2021).

We then determined the "expected range" of allele frequency change (i.e. frequency in 2021 – frequency in 1992) for each SNP as the range from the 2.5% to the 97.5% quantile of the distribution of simulated allele frequency changes in 2021. This expected range includes the expected allele frequency changes under genetic drift, sampling and the uncertainty of our model inference.

If the control SNP set contained no selected SNPs and the inference worked perfectly, ~5% of the control SNPs would fall outside this expected range and about half of the control SNPs would fall above the median of their respective expected range. In contrast, the spatial outlier SNPs were predicted to fall outside the expected range more frequently and more than half of them were predicted to fall above the median of their respective expected range, reflecting selection.

We ran the same analysis for the inversions, treating each inversion as a single locus and asking whether the arrangement frequency change exceeds the expected range. For complex inversions with three arrangements (LGC6.1/2 and LGC14.1/2), we included only the arrangement that was most common in the Wave population.

*Change in time vs space*

To compare differentiation in space and in time, we calculated the $F_{ST}$ between Crab and Wave populations based on previous studies (spatial $F_{ST}$) and the $F_{ST}$ between the skerry in 1992 and 2021 (temporal $\mathrm{F_{ST}}$) for the spatial outliers. We predicted that SNPs with a more pronounced spatial differentiation also change more dramatically over time.

For the spatial outliers obtained from Morales et al. (2019), the spatial Crab-Wave $\mathrm{F_{ST}}$ was calculated by averaging across the Crab-Wave $F_{ST}$ estimates for the locations Arsklövet, Saltö, Ramsö and Jutholmen in that study (n = 162 SNPs). These locations are all in the same archipelago as the skerry system. For the spatial outliers obtained from Westram et al. (2018), the Crab-Wave $F_{ST}$ was calculated by averaging across the Crab-Wave $F_{ST}$ estimates for all seven hybrid zones included in Westram et al. (2018) and Westram et al. (2021), which are also located in the same archipelago (n = 26 SNPs; does not include all spatial outliers from Westram et al. (2018) because of missing data for some SNPs). We then correlated spatial and temporal differentiation.

Inversion frequency estimation

We estimated the frequency of the arrangements in inversions through a clustering analysis of individuals based on PCA of SNPs in inversions. We performed a PCA on the filtered dataset, one inversion at a time including all samples, using the R package *FactoMineR* version 2.8 with default parameters. Because the first two principal components explain the largest percentage of the variance, individuals that are homozygous for each arrangement (homokaryotype) will cluster on the extremes of the PC1 vs PC2 plot, while heterozygous individuals (heterokaryotype) will cluster in-between the extremes (*25*) (Figure S10).

To identify the most likely cluster (i.e. inversion karyotype) for each individual in a population (points on the PC1 vs PC2 scatter plot), we used the implementation of k-means clustering algorithm in the R package *stats* version 4.3.0 with parameters *nstart* = 20, and centers = 3 for simple inversions and 6 for complex inversions. Considering an inversion as a multi-allele locus, we estimated the frequency of each arrangement (not to be confounded with karyotype) within a population (e.g. Skerry 1992) by simply counting the number of alleles within the clusters where the arrangement occurs divided by the total number of alleles in the population. Noteworthy, alleles in homokaryotype clusters have x2 the number of individuals, while heterokaryotype clusters have x1 the number of individuals. Last, we identified the arrangement that is more frequent in Wave than in Crab ecotype (based on merged samples from 2018 and 2021), which henceforth we will refer to as “Wave arrangement”. The arrangement frequencies are summarized in Table S6.

Simulation of inversion trajectories within the expected range of frequencies without selection

To identify the arrangements that increased in frequency most likely due to selection, we generated a neutral range expectation of arrangement frequency trajectories. Using the parameters that best describe the demographic history of the skerry (Table S7; supplementary text) and the initial arrangement frequencies in 1992, we simulated 1000 replicates of neutral trajectories of each inversion under drift and migration. In every generation $T$ (out of 48 until 2021, the last sampling time point), we randomly sampled $N_T$ alleles (population growth based on $r$ and $K$ from the best model) from a binomial distribution with a probability equal to the frequency of the Wave arrangement in $T_{-1}$plus the proportional contribution of migrants $M$. We determined the neutral range for each inversion as the range from the 2.5% to the 97.5% quantile of the distribution of simulated arrangement frequencies in 2021. Thus, arrangements whose frequency in 2021 exceed the neutral range are likely be under selection. Figure S11 shows the observed and simulated trajectories of simple and complex inversions.

**Supplementary Text**

Results of selection estimates based on phenotypes

Using $V_g$-*crab*, estimates of the strength of stabilising selection ($V_s/V_p$) ranged from 1.65 to 7.84 on the assumption of one generation per year and from 3.80 to 25.3 on the assumption of two generations per year, excluding the aperture position trait (r0) for which all of the change on the skerry could be accounted for by plasticity (Table S2). Weaker selection was inferred using the larger estimate of genetic variance ($V_g$-*site*). The strongest selection was on width growth (gw), for which the skerry phenotype was already close to the Wave reference phenotype in 2002. Other traits were approaching the reference phenotype in the final two samples (2018 and 2021). The estimated strengths of stabilising selection are in the range observed in surveys of phenotypic selection in nature, where $V_s/V_p$ is typically ~5 (*31*, *32*)However, this translates into strong selection on the Crab phenotype in the skerry environment because of its distance from the optimum, with some estimated fitness reductions >90% under the assumption of one generation per year and using the $V_g$ estimate for the Crab populations (Table S2).

Results of the demographic inference

*Estimated demographic parameters*

The most likely combination of parameters from the grid given the difference in log likelihood of that combination vs the maximum is shown in Table S7. The means and support limits for the different parameters based on interpolation are shown in Table S8. Note that the parameters referring to counts of individuals reflect haploid individuals; the diploid value would be half the estimates shown here.

*Expected range*

For both control and spatial outlier SNPs, the proportions of SNPs inside and outside the expected range, and above and below the median value from 1,000 replicate simulations are shown in Table S9 and illustrated in Figure S12. For the control SNPs, we found that 8% of SNPs were outside the expected range, i.e. slightly more than the 5% expected if the model fitted perfectly and the SNPs were only affected by drift. The discrepancy is unsurprising given that some SNPs may be affected by linked selection and that it is unlikely that our simple model perfectly captures the history of the skerry. The pattern is more or less symmetrical (almost as much change away from Wave as towards Wave).

For spatial outliers, we found a similar proportion outside the expected range (9%); however, there was a clear asymmetry, with almost all SNPs outside the neutral range showing more change towards the Wave allele frequency than expected and with 71% of SNPs showing more change towards Wave than the median change under neutrality. These results indicate that at least a large subset of the spatial outlier SNPs are affected by selection. However, it is difficult to say which exact SNPs are affected by selection, as the extent of drift and / or the model uncertainty are too high for most SNPs to show statistically significant changes (i.e. be outside the expected range).

For the inversions, we found that all 13 inversions included in the analysis showed an arrangement frequency change above the median of the respective expected range, with 4 (31%) showing a change outside the expected range. These results strongly indicate that multiple inversions are affected by selection on the skerry.

### Supplementary Figures

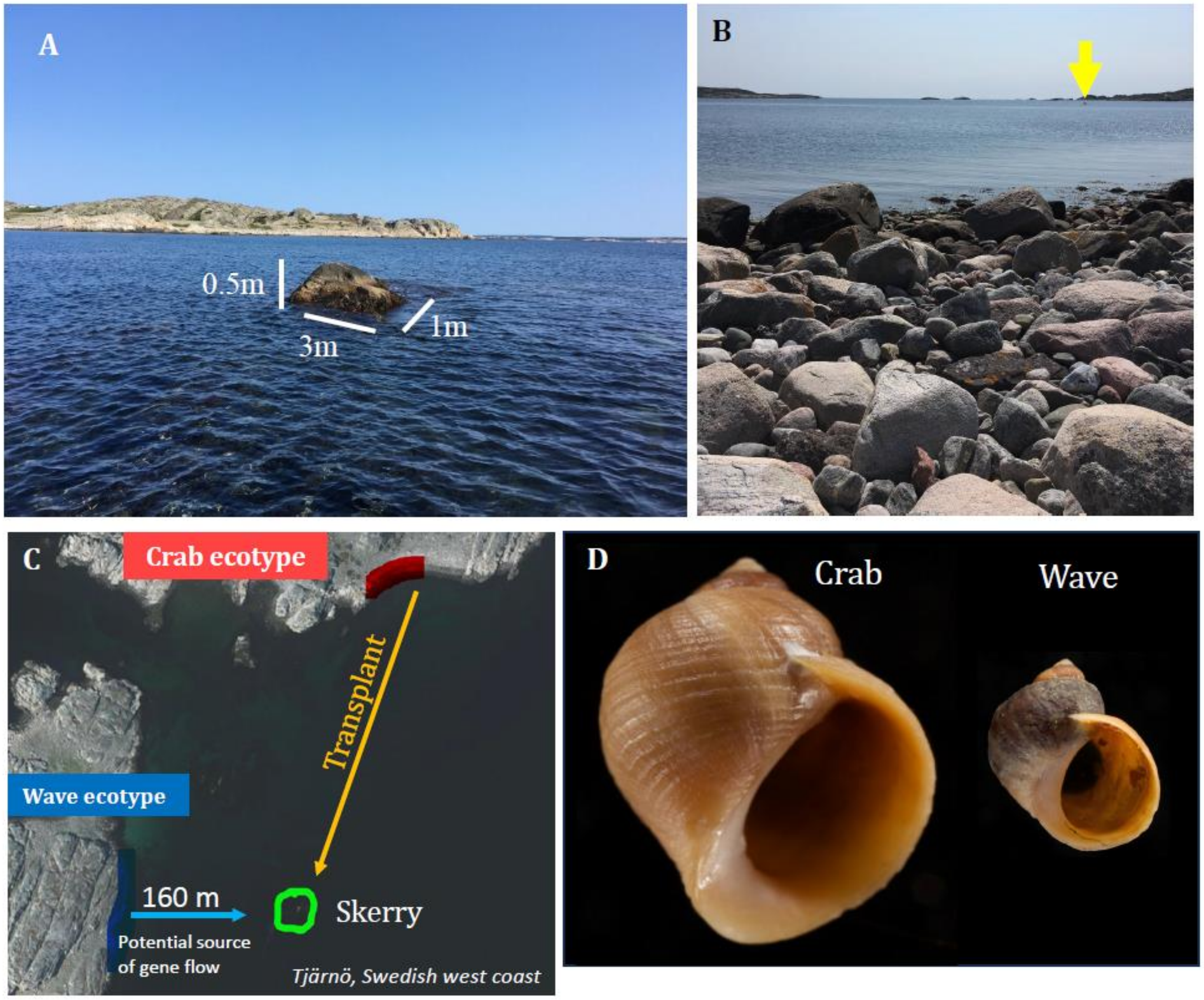


**Figure S1. The skerry area. (A)** Photography and dimensions of the skerry, the target of the transplanted snails from a Crab ecotype. **(B)** A photography of the skerry (yellow arrow) taken from the shore of the donor Crab ecotype. **(C)** A satellite view of the skerry, the donor Crab ecotype, and the neighbouring Wave ecotype. **(D)** Photography of two sample shells from the Crab ecotype and the Wave ecotype.

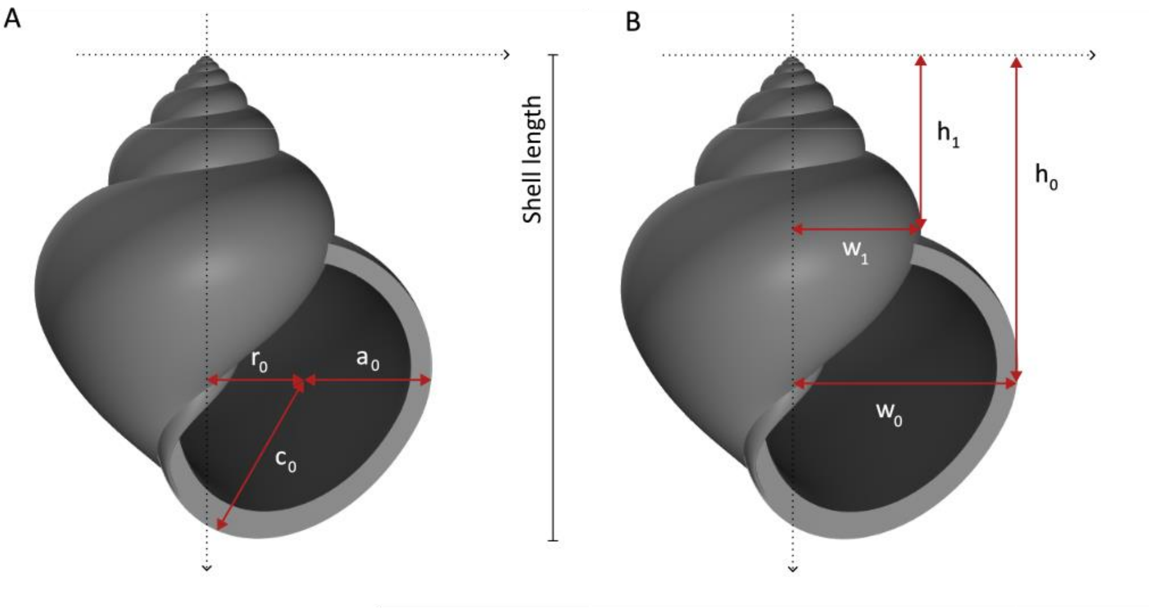


**Figure S2: Description of the shape parameters used in the analysis.** All measurements are relative the total shell length, to remove the size component.

$a_0$ : Size of the (circular, upper part of the) aperture.

$r_0$ : Radial position of the aperture reference point (centre of the circular part).

$h_0$ : Vertical position of the aperture reference point

c : Shape of the aperture, measuring elliptical eccentricity of the lower part of the aperture ($\boldsymbol{c} = \frac{\boldsymbol{c_0}}{\boldsymbol{a_0}}$)

$g_w$: Width growth between consecutive whorls ($\boldsymbol{g_w} = \frac{\boldsymbol{w_0}}{\boldsymbol{w_1}}$).

$g_h$: Height growth between consecutive whorls ($\boldsymbol{g_h} = \frac{\boldsymbol{h_0}}{\boldsymbol{h_1}}$).

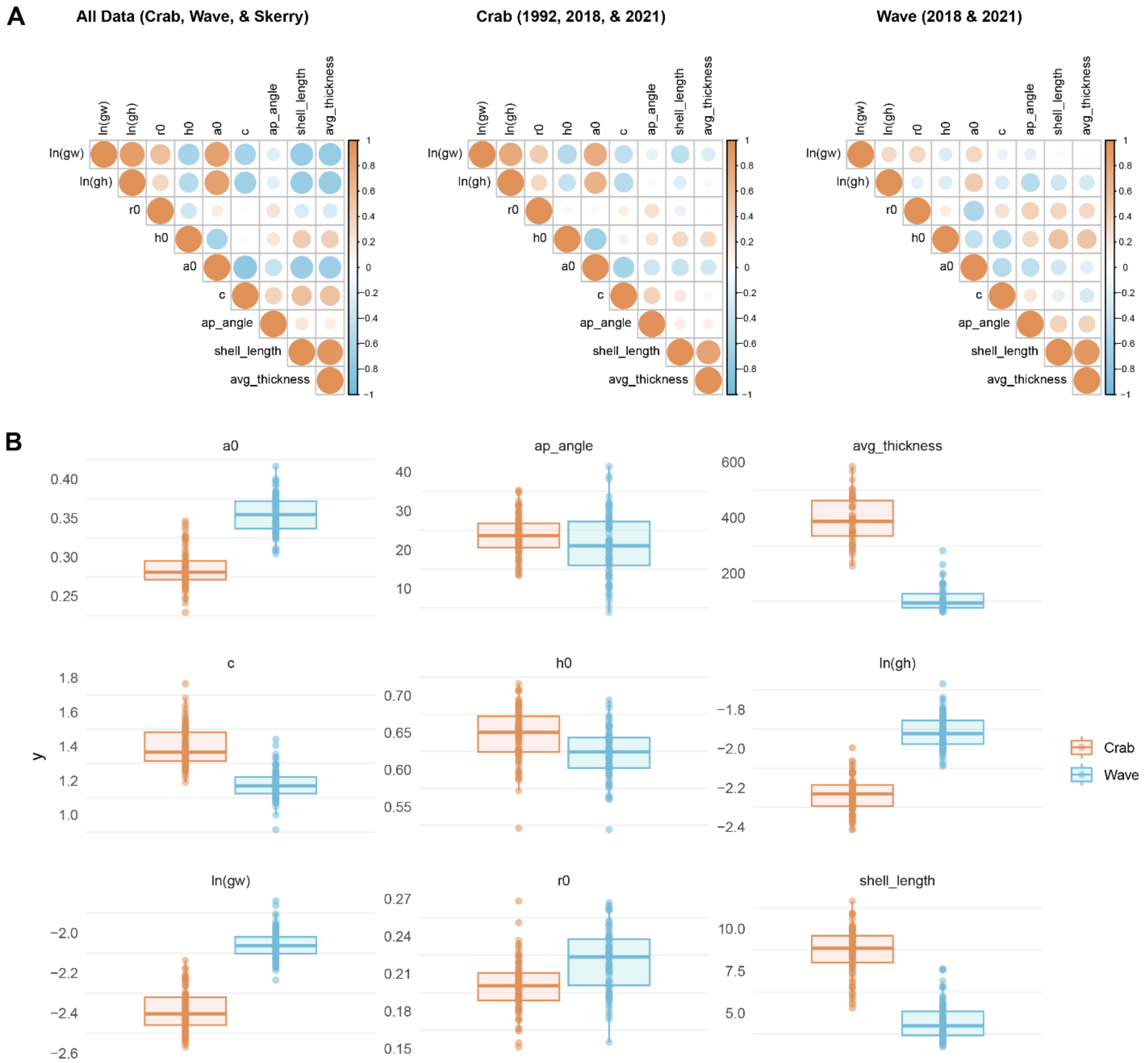


**Figure S3. Comparison between quantitative traits. (A)** Correlation plots of different quantitative traits in all populations, the Crab ecotype, and the neighbouring Wave ecotype. The color scale represents the correlation coefficient between each pair of traits. **(B)** Boxplots of pairs of diagnostic traits in the Crab ecotype and neighbouring Wave ecotype.

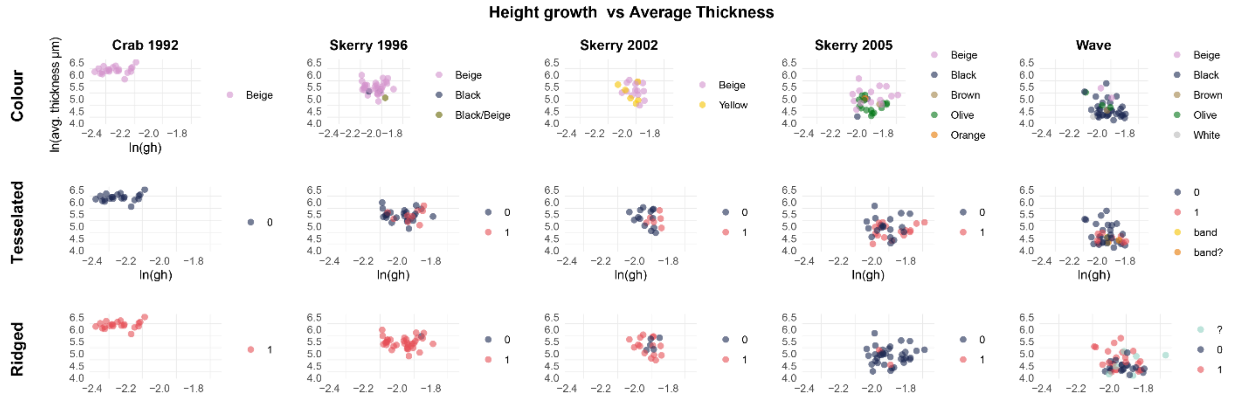
Height growth vs Average Thickness
Crab 1992
Skerry 1996
Skerry 2002
Skerry 2005
Wave
Colour
Tesselated
Ridged
ln(avg. thickness μm)
ln(gh)
Beige
Black
Black/Beige
Yellow
Brown
Olive
Orange
White
0
1
band
band?
?

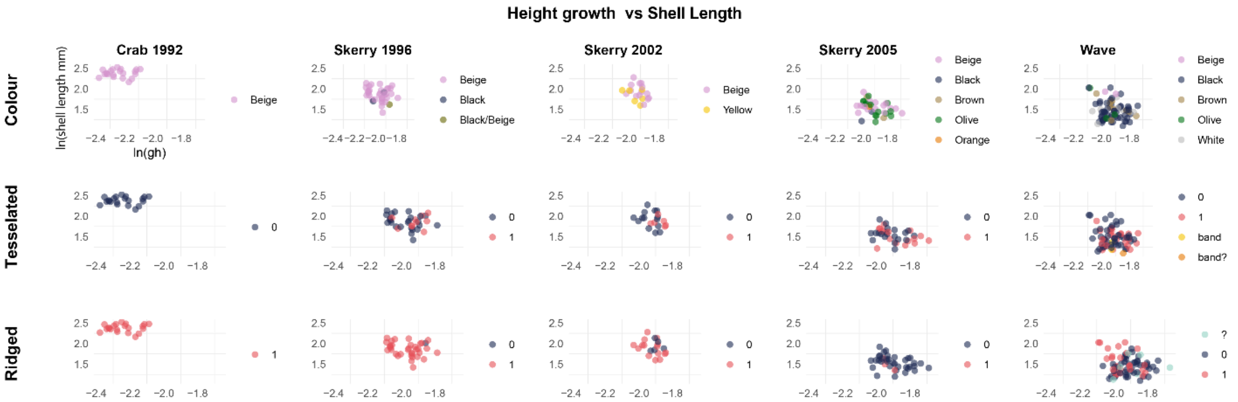
Height growth vs Shell Length
Crab 1992
Skerry 1996
Skerry 2002
Skerry 2005
Wave
Colour
Tesselated
Ridged
ln(shell length mm)
ln(gh)
Beige
Black
Black/Beige
Yellow
Brown
Olive
Orange
White
0
1
band
band?
?

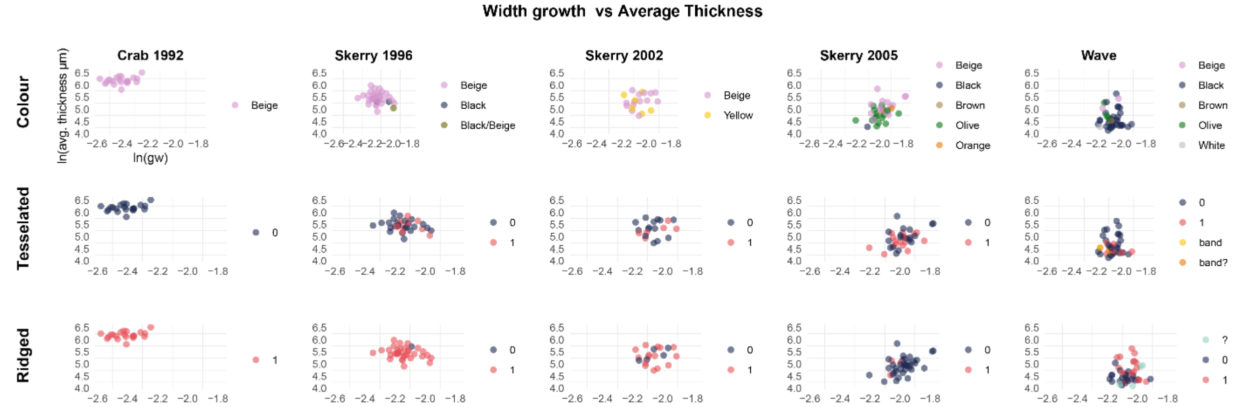
Width growth vs Average Thickness
Crab 1992
Skerry 1996
Skerry 2002
Skerry 2005
Wave
Colour
Tesselated
Ridged
ln(avg. thickness μm)
ln(gw)
Beige
Black
Black/Beige
Yellow
Brown
Olive
Orange
White
0
1
band
band?
?

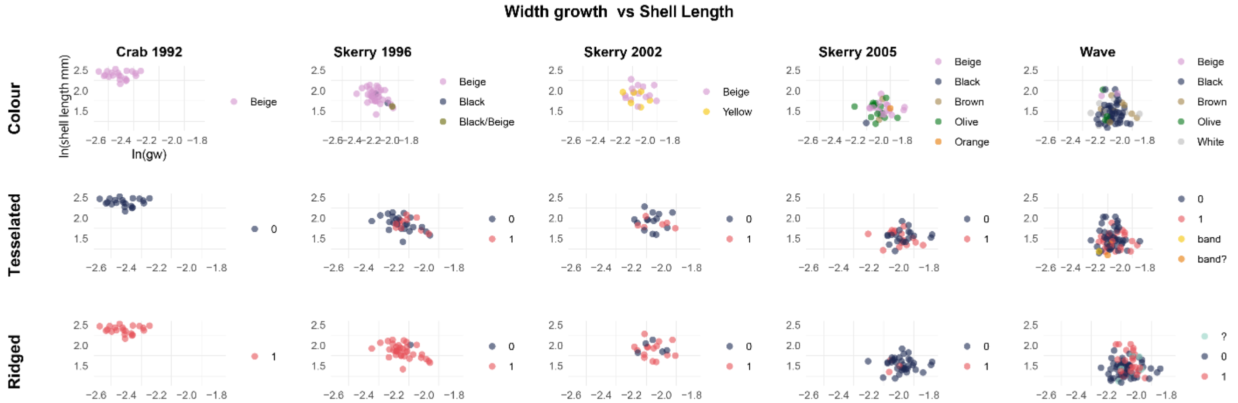
Width growth vs Shell Length
Crab 1992
Skerry 1996
Skerry 2002
Skerry 2005
Wave
Colour
Tesselated
Ridged
ln(shell length mm)
ln(gw)
Beige
Black
Black/Beige
Yellow
Brown
Olive
Orange
White
0
1
band
band?
?

**Figure S4. Scatter plot of pairs of uncorrelated quantitative traits for the first few years after the introduction.** Rows are three groups of qualitative traits. Samples are coloured by the different categories of a qualitative trait. All four quantitative traits (average thickness, average shell length, height growth, and width growth) were transformed to natural logarithm (ln).

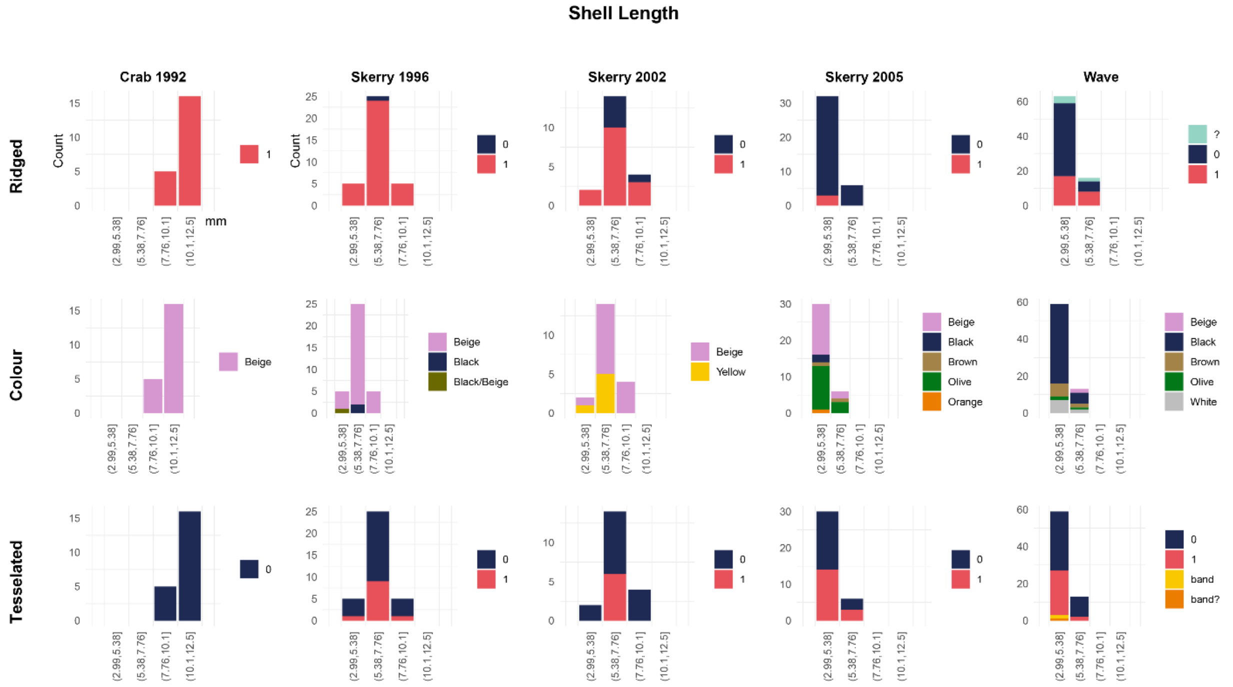
Shell Length
Crab 1992
Skerry 1996
Skerry 2002
Skerry 2005
Wave
Ridged
Colour
Tesselated
Count
mm
Beige
Black
Black/Beige
Yellow
Brown
Olive
Orange
White
band
band?

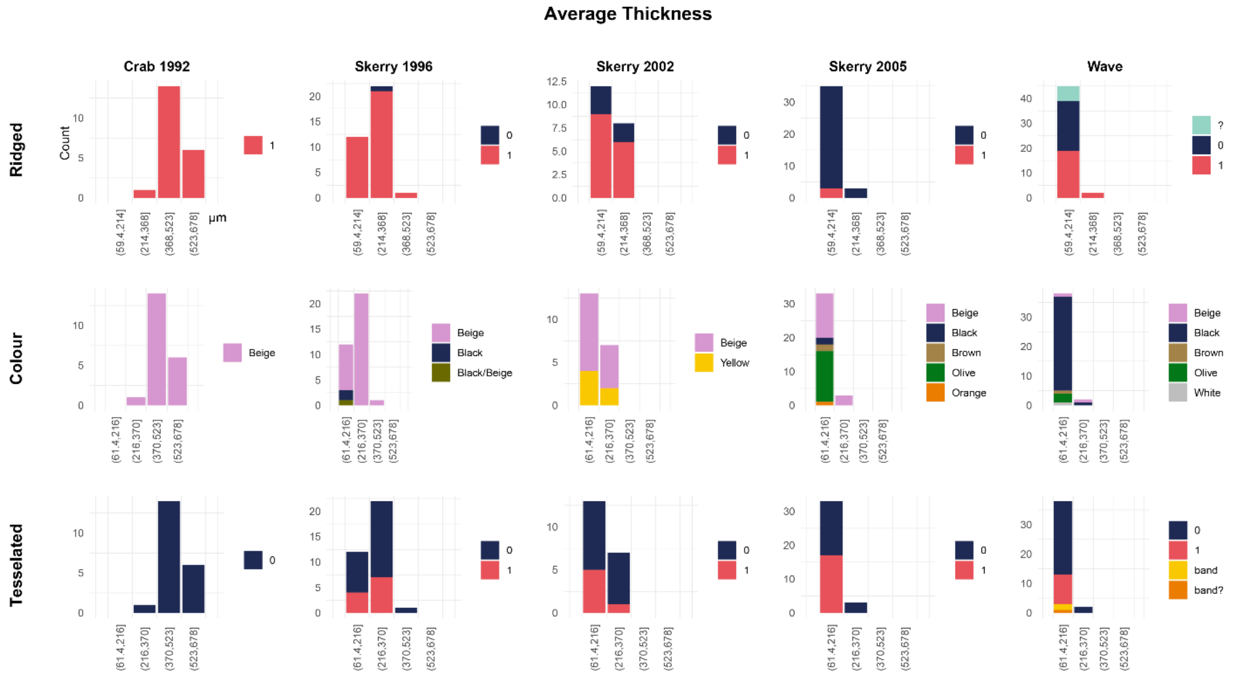
Average Thickness
Crab 1992
Skerry 1996
Skerry 2002
Skerry 2005
Wave
Ridged
Colour
Tesselated
Count
µm
Beige
Black
Black/Beige
Yellow
Brown
Olive
Orange
White
band
band?

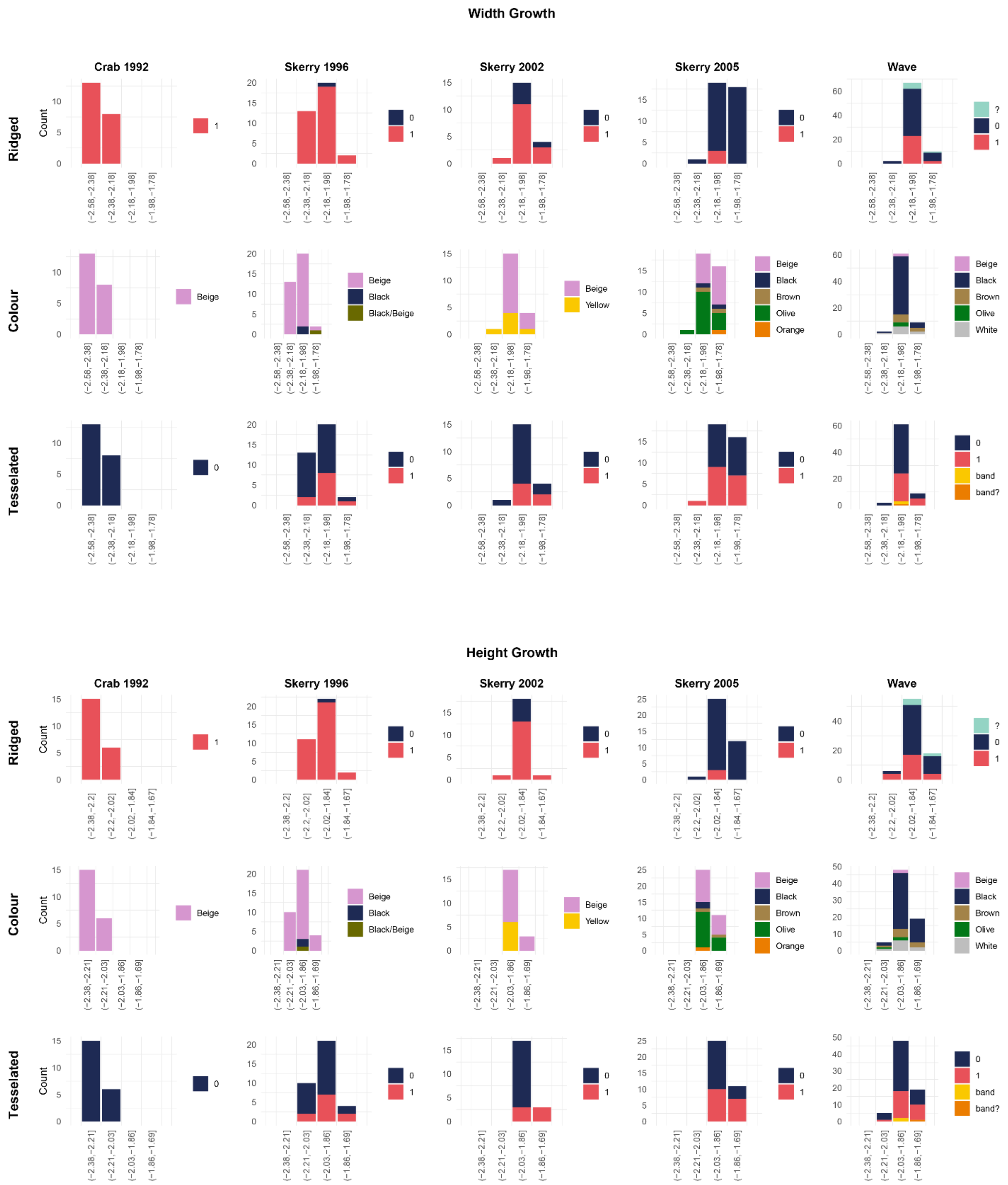


**Figure S5. Bar plots of four diagnostic traits (shell length, average thickness, width growth, and height growth) the first few years after the introduction.** Rows are three groups of qualitative traits. Bars were plotted in four bins, and coloured by the counts in different categories of a qualitative trait within each bin.

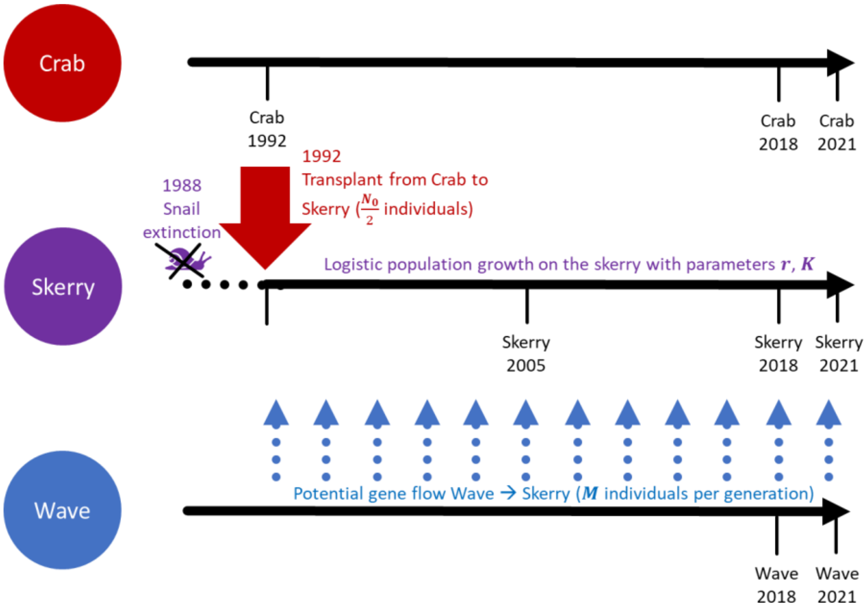


**Figure S6: Schematic depiction of the experiment, with parameters of the demographic model indicated in bold.** The black arrows represent time; sampling times are indicated below. Events important for the experiment are highlighted in colour.

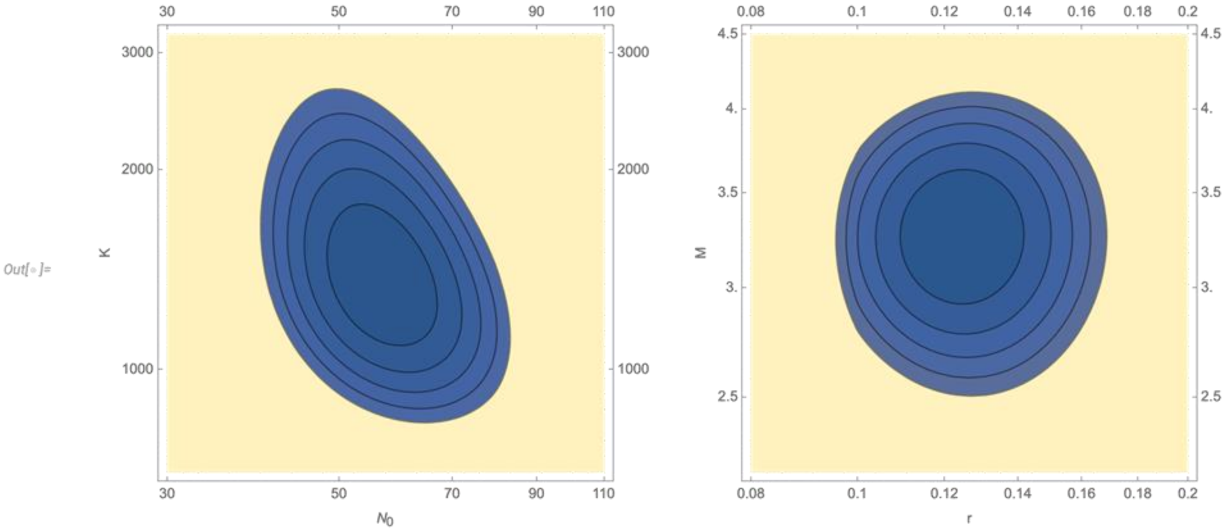


**Figure S7. Contours of log likelihood, with spacing 1, as a function of $N_0$, K (left) or r, M (right).** In each plot, the other two parameters are fixed at their MLE (left: r=0.12, M=3.26; right: $N_0$=55.6, K=1335). For two degrees of freedom, a loss of log likelihood of 3 corresponds to $\chi_2^2$, or P=5%. Thus, three contours down corresponds to 95% confidence intervals.

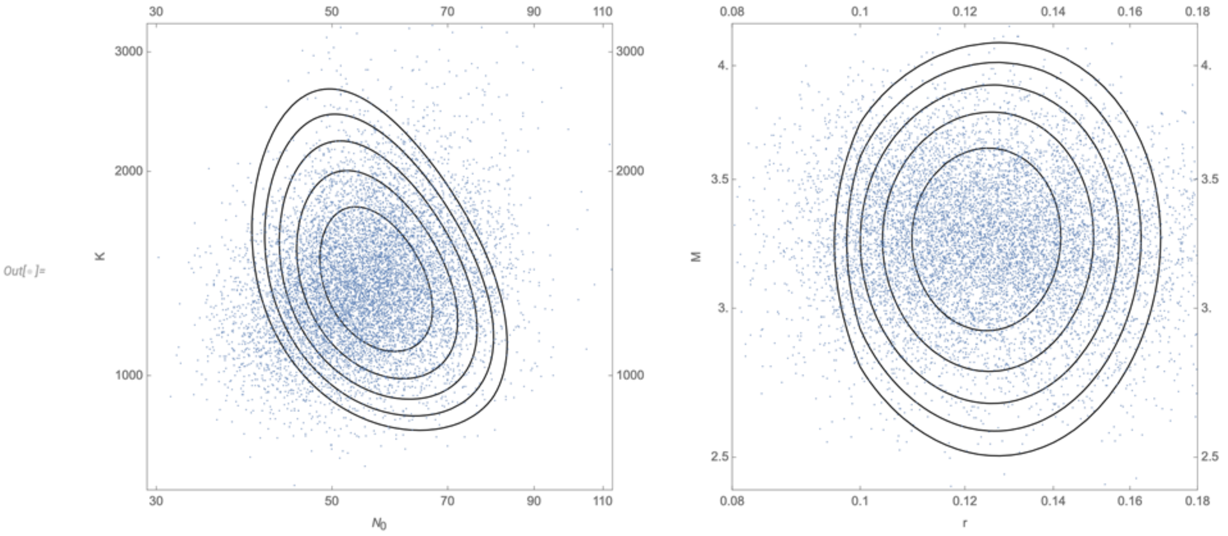


**Figure S8. Posterior distribution (points) superimposed on contours of log likelihood (spacing 1).** Left: $N_0$ vs K. right: r vs. M. Contours on the left plot fix f=2, M=3.26, r=0.12; the right plot fixes $N_0$=55.6, K=1335. Note that these distributions are not quite the same: the posterior distribution averages over the posterior distribution of the other two parameters, whereas the contours show the log likelihood with the other two parameters fixed at their MLE.

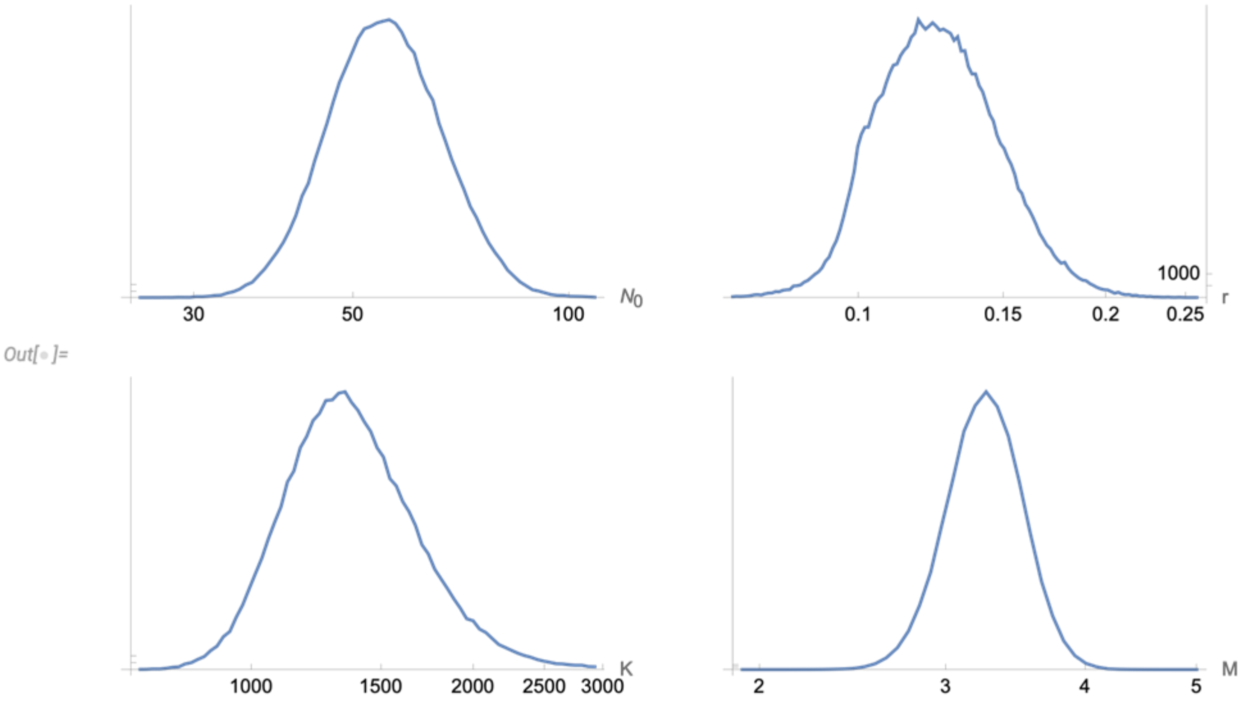


**Figure S9. Posterior distribution for the four parameters, based on 500,000 random draws, using the Metropolis algorithm.**

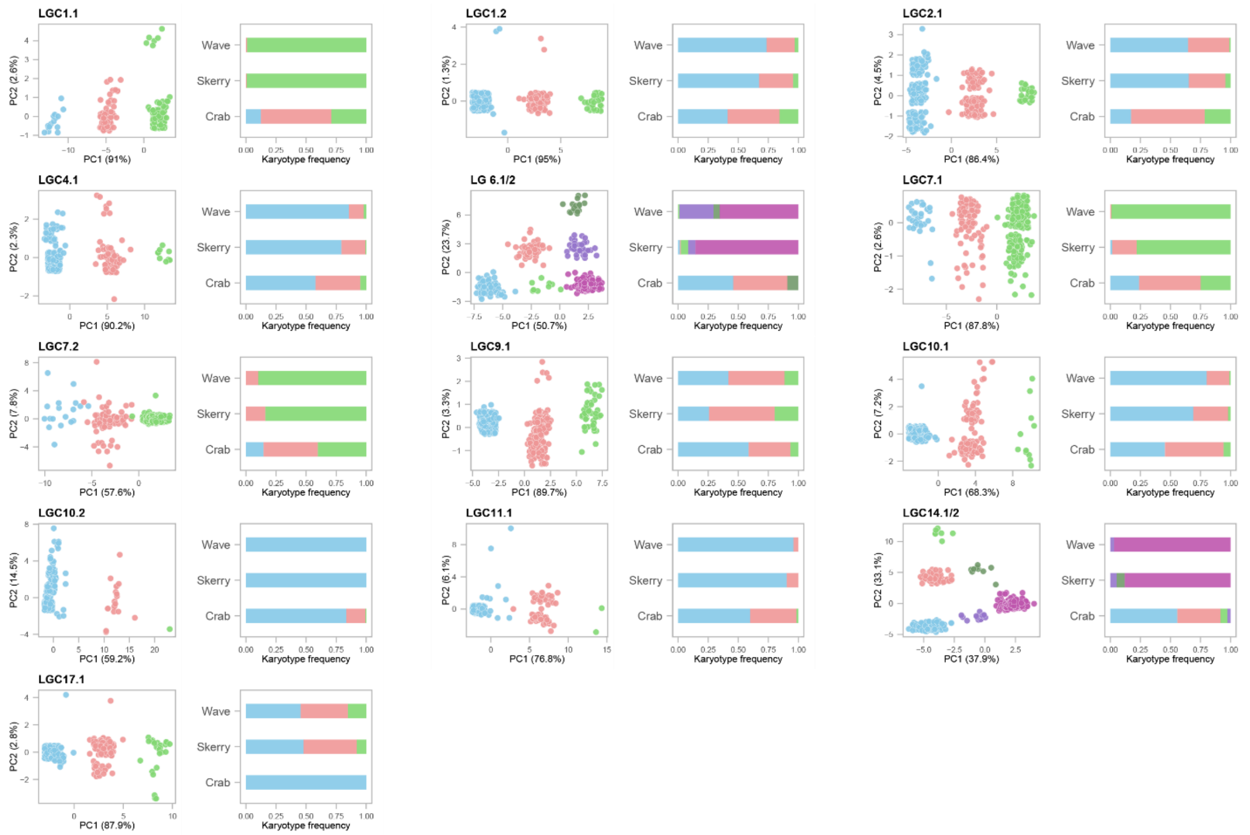


**Figure S10. Clustering pattern of PC1 vs PC2 for both simple and complex inversions**. The bar plots next to each scatterplot shows the frequency of the different karyotypes for a particular inversion in the skerry, Crab (1992, 2018, and 2021) and Wave (2018 and 2021) ecotype populations. The points were coloured based on the most likely partitioning by a k-means algorithm. The colours in the bars correspond to the colours of the clusters in the scatterplot.

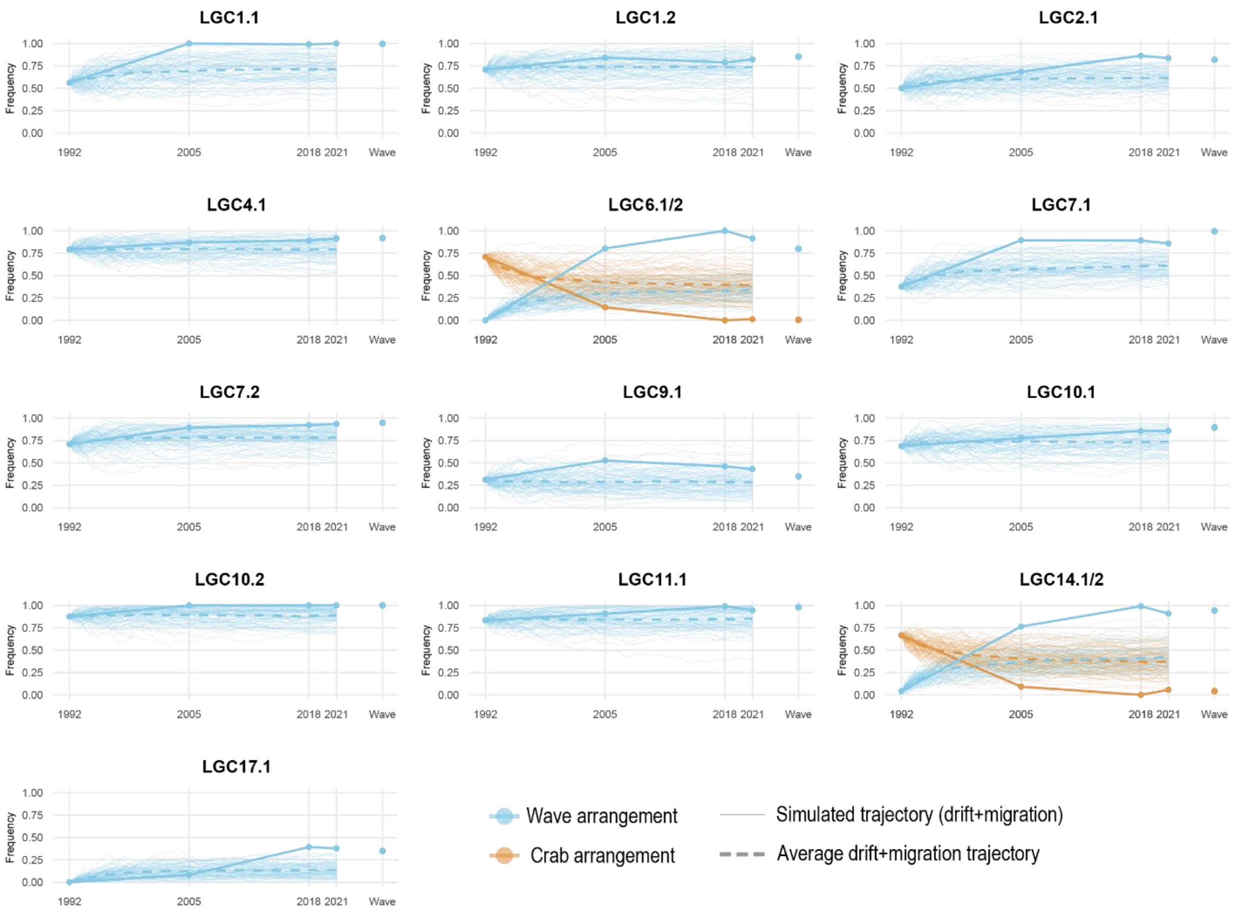


**Figure S11 Frequency trajectories of the Wave arrangement**. In blue, trajectories of the Wave arrangement. In orange, trajectories of the Crab arrangement in complex inversions (two out of three arrangements). To facilitate visualization, the figures include 100 replicates of simulated trajectories for each arrangement (thin and pale lines). However, the neutral range was generated from 1000 simulated trajectories for each arrangement.

**A**

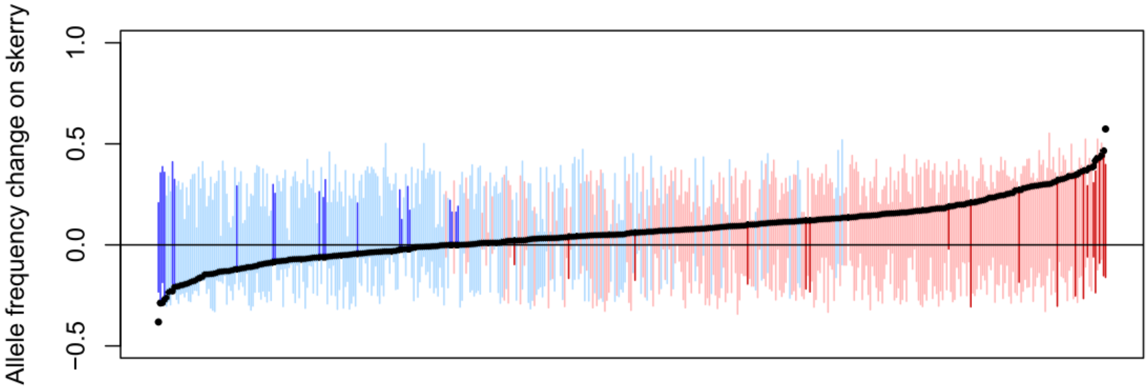


**B**

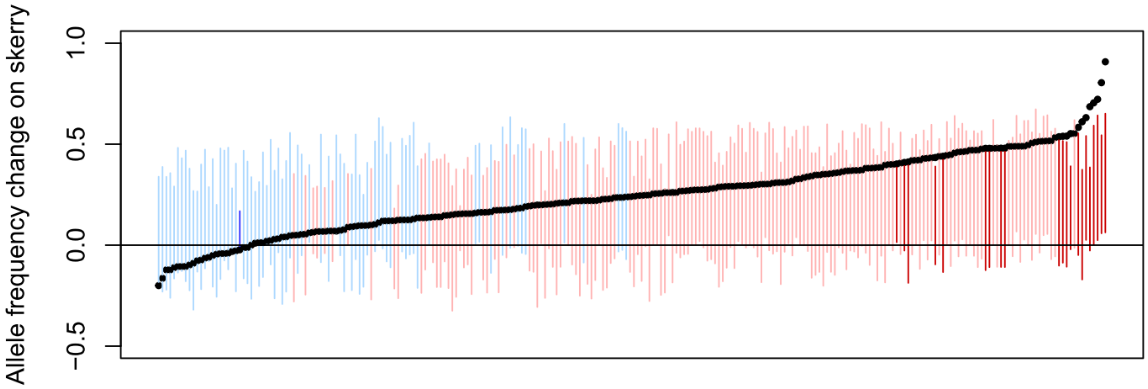


**C**

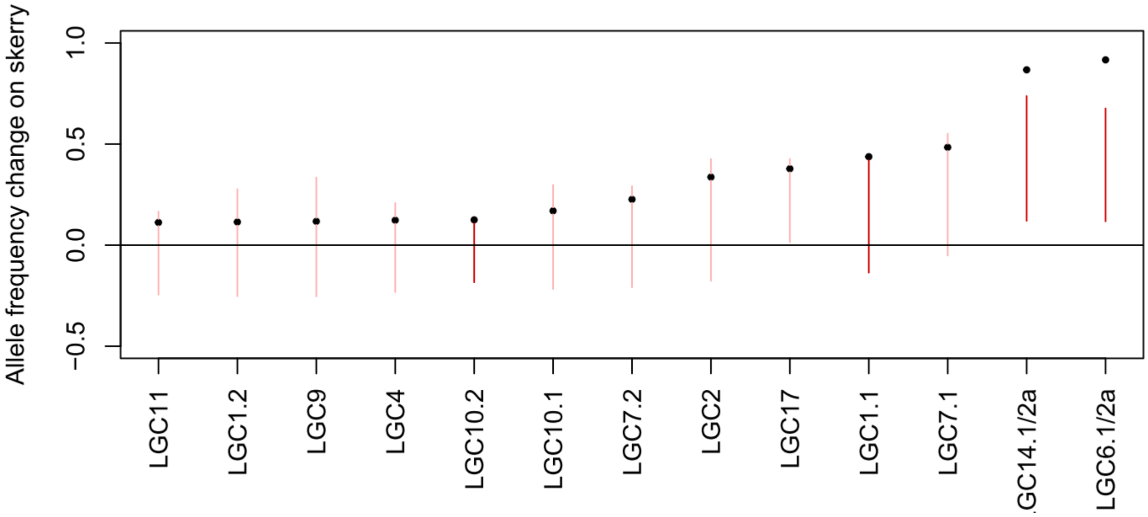


**Figure S12. Expected ranges (coloured bars) and observations (black circles) for the allele frequency change on the skerry (i.e. the allele frequency difference between the skerry population in 2021 and the Crab donor population in 1992).** Positive allele frequency change indicates change towards the allele more common in Wave, while negative change indicates that the frequency of the allele more common in Wave decreased. SNPs are sorted along the x-axis by the extent of allele frequency change. A) control SNPs; B) spatial outlier SNPs; C) inversions. Dark blue: observation outside expected range and below median; light blue: observation inside expected range and below median; light red: observation inside expected range and above median; dark red: observation outside expected range and above median. For the inversions, inversion IDs are indicated along the x-axis. For complex inversions with three arrangements, only the arrangement more common in Wave than in Crab is shown.

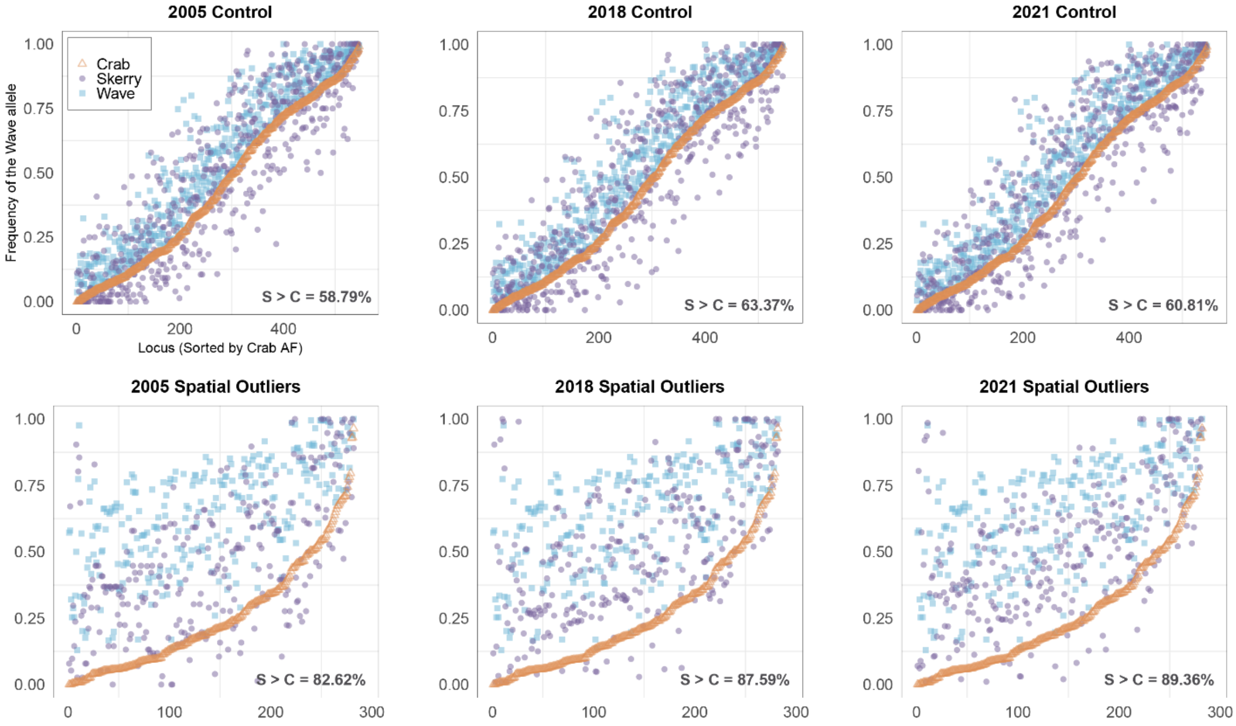


**Figure S13. Frequency of the Wave allele in the skerry and reference populations.** "Wave" allele refers to the allele that is more common in the Wave (merged samples from 2018 and 2021) than in the Crab (merged samples from 1992, 2018, and 2021) ecotype populations. We sorted the loci in ascending order, according to the allele frequency (AF) in Crab. In all sampled years, the frequency of the Wave allele in skerry was more commonly observed above the frequency found in Crab (the orange middle pattern is the result of adjacent SNPs sorted by AF in crab). As expected due to their role in ecotype divergence, spatial outliers (bottom row) changed more frequently towards the Wave frequency compared to control loci (top row).

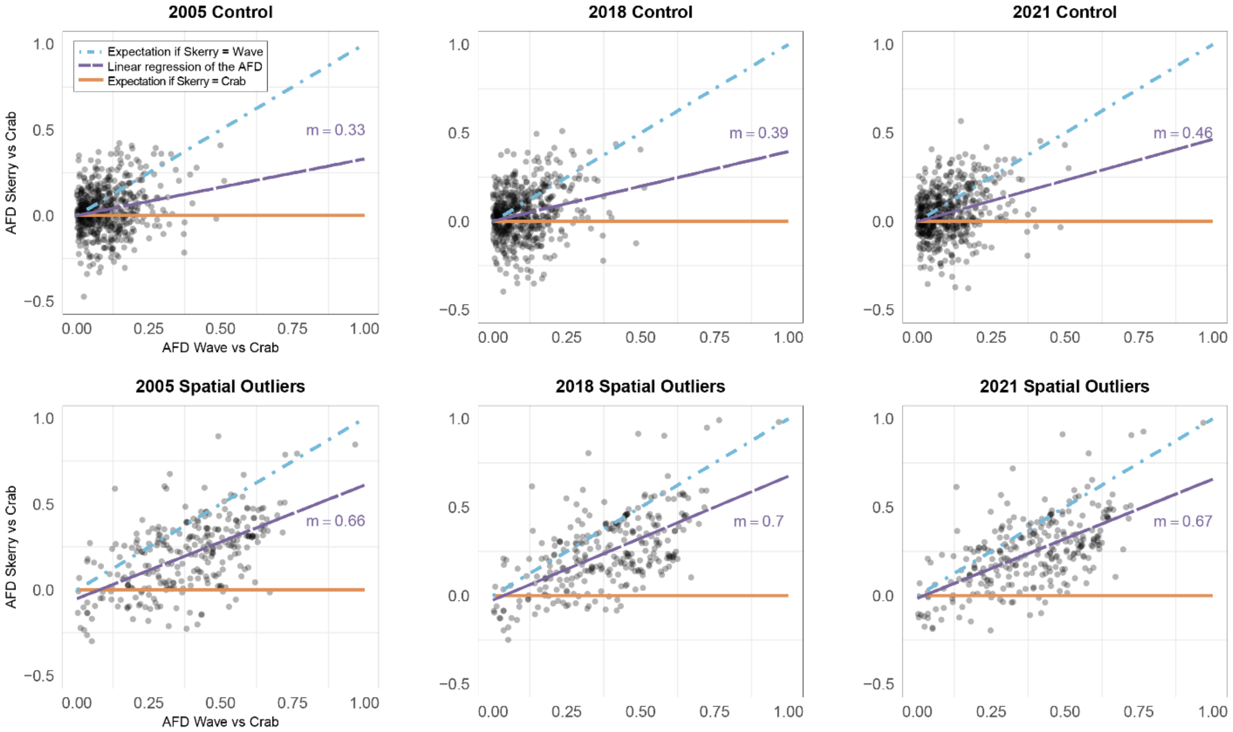


**Figure S14. Directional allele frequency change in the skerry in three sampling years.** Allele frequency difference (AFD) estimates are based on the frequency of the "Wave" allele (allele that is more common in Wave than in Crab). The allele frequencies of Wave included samples from 2018 and 2021. Likewise, the allele frequencies of Crab included samples from 1992, 2018, and 2021.The slope (m) of a linear regression (dashed line) indicates a directional allele frequency change towards Wave in spatial outliers (bottom row) but not in control loci (top row), as predicted. The dashed blue line represents the expectation if the allele frequencies (AFs) in the skerry population were identical to those in the Wave ecotype. The solid orange line represents the expectation if the AFs in the skerry population were identical to those in the Crab ecotype.

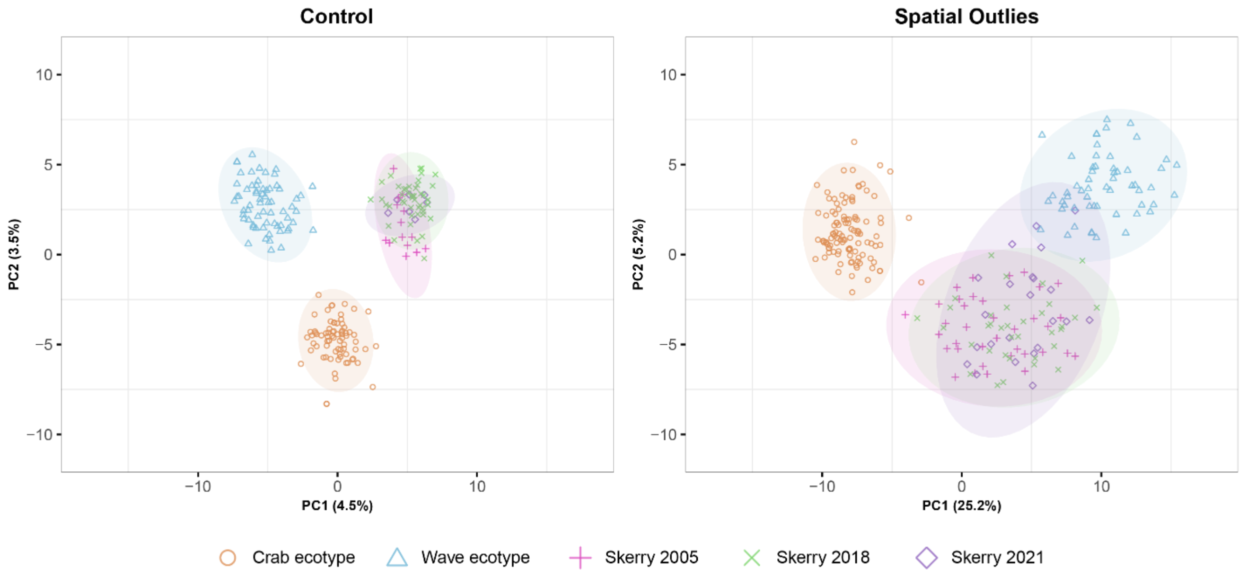


**Figure S15. PCA on collinear loci.** Crab ecotype (Donor Crab) includes the samples from 1992, 2018, and 2021. Wave ecotype includes the samples from 2018 and 2021. The clustering pattern of control loci suggests a strong effect of drift that separates genetically the skerry from the donor population (Crab) on both axes. On the contrary, for candidate outliers, the skerry samples cluster closer to the Wave ecotype on PC1 (the axis that explains a larger proportion of the total variance) likely due to a strong effect of selection. This plot also shows that most change on the skerry happened early in the experiment, as there is little change from 2005 to 2021.

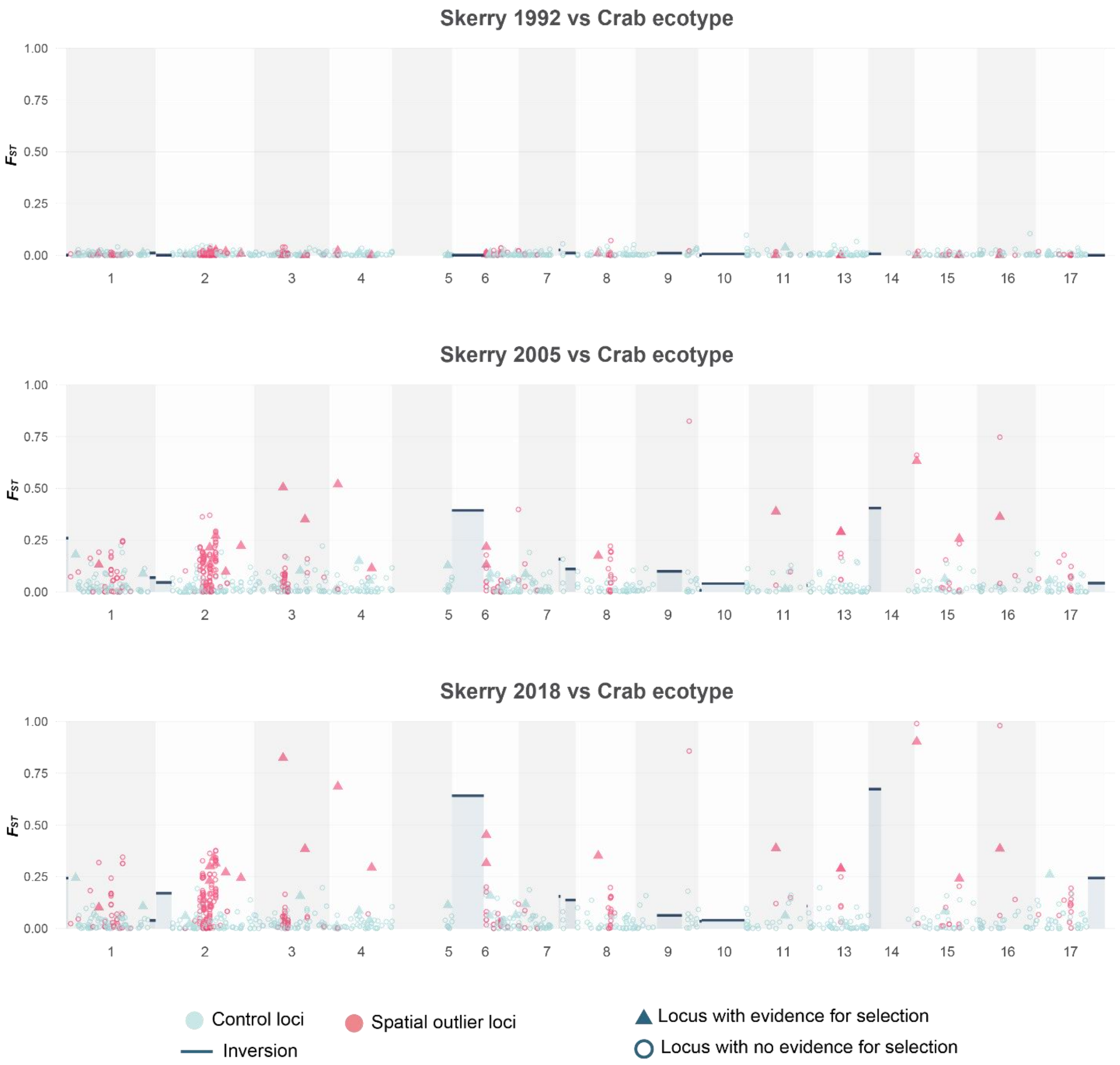


**Figure S16. Genome-wide $F_{ST}$ in the skerry versus the average crab ecotype in three different years.** Circles and triangles represent individual SNPs in the collinear genome. Inversions are represented by rectangular blue-grey fields with black bars at top indicating $F_{ST}$ value.

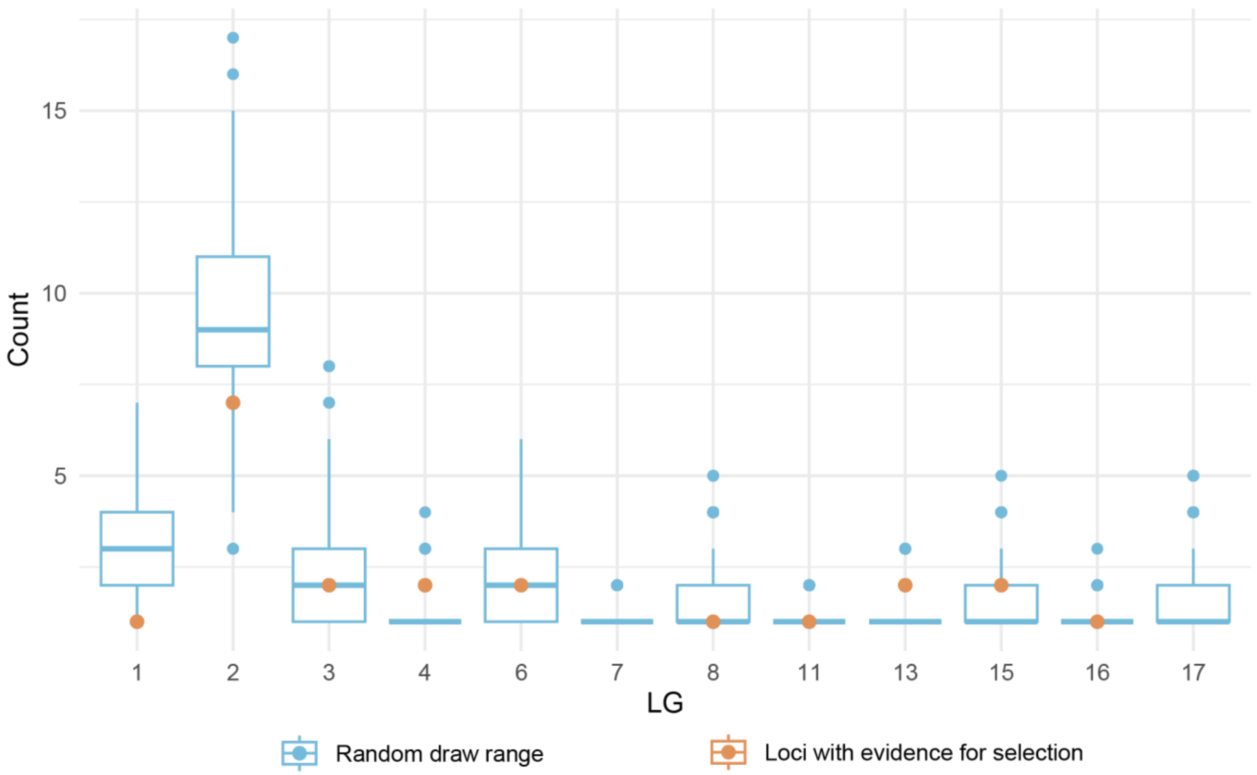


**Figure S17. Count of spatial outliers with evidence for selection (orange) with respect to the expectation based on SNP content in each chromosome (blue).** We randomly sampled 21 spatial outliers (spatial outliers with evidence for selection based on the expected range under neutrality) along the collinear genome 1000 times. The random draws, which are the chance expectation, are plotted as blue boxes. Only linkage groups with more than one spatial outliers were included.

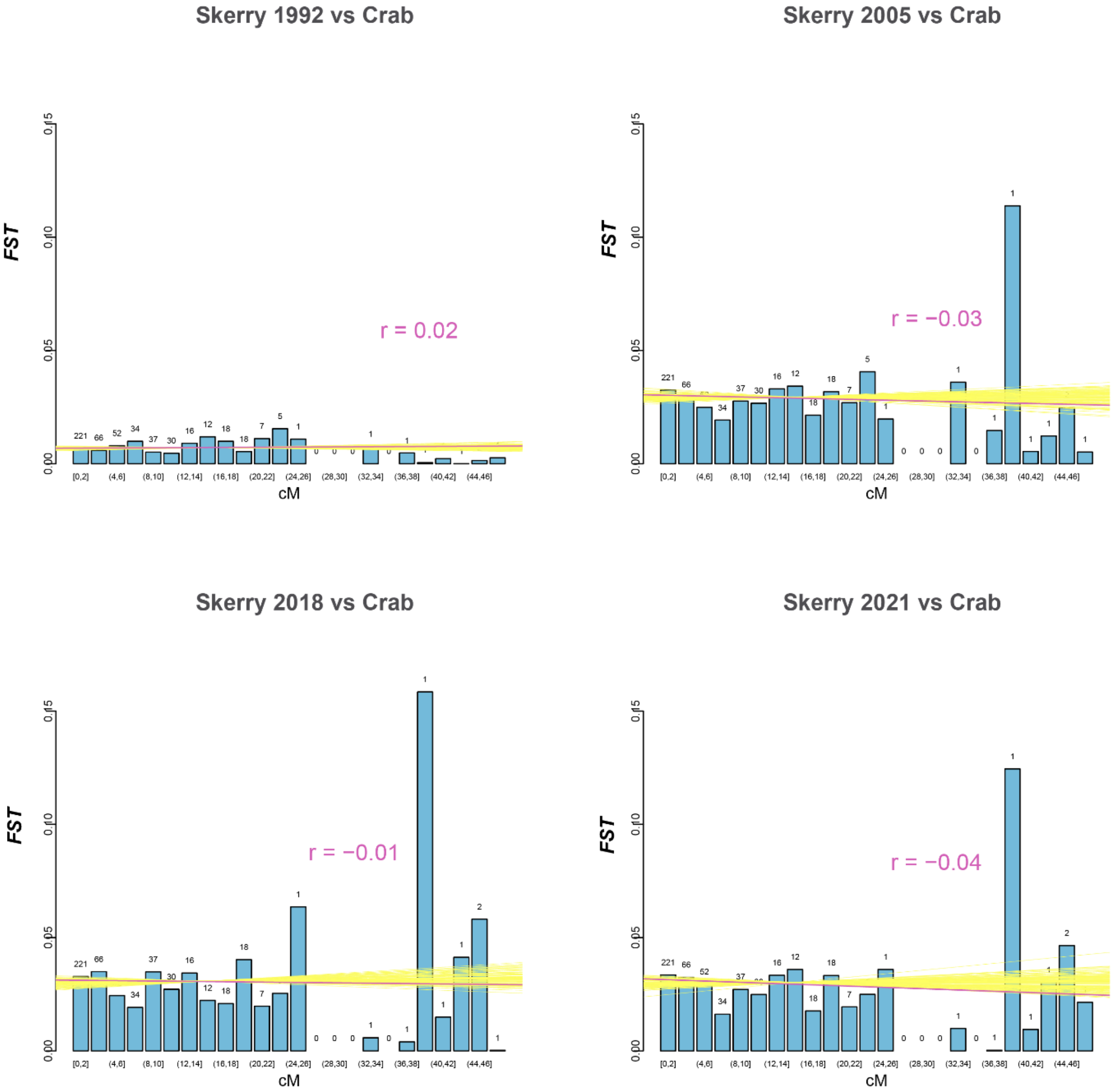


**Figure S18. Correlation between genomic distance and genetic differentiation (*$F_{ST}$*).** The barcharts show the $F_{ST}$ of control loci in the Skerry from different years vs the average Crab ecotype (2018+2021). The x axis is the distance in cM of each control locus to the nearest spatial outlier. The purple line is the $F_{ST}$-to-distance linear regression of the empirical data, accompanied by the correlation coefficient (r). In 2005, 2018, and 2021, the $F_{ST}$ value is weakly negatively correlated to the distance to the nearest candidate outlier locus. To test whether this pattern can be generated by chance, we shifted randomly the coordinates of control loci and estimated both the linear regression and the correlation coefficient. The yellow lines are multiple simulations of $F_{ST}$-to-distance linear regression. The correlation observed in empirical data is always within the range of shuffled loci. Thus, there is no evidence of strong hitch-hiking effects, although, it does not rule out genome-wide effects in the early generations, when selection is very strong.

### Supplementary Tables

**Table S1. Number of markers from each category**

| Marker type | Marker number |
|---|---|
| Baypass 2* | 28 |
| Baypass 3* | 32 |
| $F_{ST}$ outlier in 1 population* | 8 |
| $F_{ST}$ outlier in 2 population* | 22 |
| $F_{ST}$ outlier in 3 population* | 33 |
| $F_{ST}$ outlier in 4 population* | 33 |
| $F_{ST}$ outlier in 5 population* | 44 |
| Clinal marker* | 102 |
| Neutral marker (control) | 565 |
| Markers in inversions | 225 |

*Spatial outliers

**Table S2. Phenotypic analysis.**

Supplementary Table Phenotypic Analysis.xlsx (External)

**Table S3: Genotyping data used in the inference.** "Usage" indicates the corresponding variable in the model and "diploid sample size" gives the number of diploid individuals to which the data set was downsampled in order to avoid missing data.

| **Sample** | **Usage** | **Diploid sample size** |
|---|---|---|
| Crab donor 1992 | $k_{1992}$, observed allele count on the skerry 1992 | 18 |
| Skerry 2005 | $k_{2005}$, observed allele count on the skerry 2005 | 31 |
| Skerry 2018 | $k_{2018}$, observed allele count on the skerry 2018 | 45 |
| Skerry 2021 | $k_{2021}$, observed allele count on the skerry 2021 | 34 |
| Wave 2018 and 2021 | Wave allele frequency $p_W$, source of gene flow | 77 |

**Table S4**: **Parameters of the demographic model.**

| Parameter | Description | Values |
|---|---|---|
| $N_0$ | Haploid starting population size on the skerry | 20, 40, 80, 160, 320 |
| $r$ | Growth rate in logistic population growth model | 0.025, 0.05, 0.1, 0.2, 0.4 |
| $K$ | Haploid carrying capacity in logistic population growth model | 250, 500, 1000, 2000, 4000 |
| $M$ | Number of haploid migrants per generation | 0, 1, 2, 4, 8 |
| $f$ | Number of generations per year | 4/3, 5/3, 2 |

**Table S5. The mean and 95% limits of the posterior distributions, based on 500,000 random draws, using the Metropolis algorithm.**

| | mean | 95% limits |
|---|---|---|
| $N_0$ | 55.4 | {39.2, 79.1} |
| **r** | 0.12 | {0.092, 0.176} |
| **K** | 1371 | {950, 2168} |
| **M** | 3.25 | {2.77, 3.77} |

**Table S6: Frequencies of the Wave arrangement in the skerry and reference populations.** Complex inversions (LGC6.1/2 and LGC14.1/2) have two rows in the table, corresponding to the Wave and the "Crab" arrangement in that order.

| Inv_ID | pC1992 | pS2005 | pS2018 | pS2021 | pW2018+2021 | pC2018+2021 |
|---|---|---|---|---|---|---|
| LGC1.1 | 0.5625 | 1 | 0.990196 | 1 | 0.994949495 | 0.587628866 |
| LGC1.2 | 0.708333 | 0.842105 | 0.788462 | 0.822917 | 0.852941176 | 0.608247423 |
| LGC2.1 | 0.5 | 0.684211 | 0.862745 | 0.836735 | 0.818181818 | 0.474226804 |
| LGC4.1 | 0.791667 | 0.868421 | 0.892157 | 0.914894 | 0.918367347 | 0.757731959 |
| LGC6.1/2 | 0 | 0.802632 | 1 | 0.916667 | 0.798913043 | 0.677083333 |
| LGC6.1/2 | 0.708333 | 0.144737 | 0 | 0.02381 | 0.005434783 | 0 |
| LGC7.1 | 0.375 | 0.894737 | 0.892157 | 0.858974 | 0.994680851 | 0.536082474 |
| LGC7.2 | 0.708333 | 0.894737 | 0.921569 | 0.934783 | 0.947916667 | 0.607142857 |
| LGC9.1 | 0.3125 | 0.526316 | 0.460784 | 0.430233 | 0.348958333 | 0.221649485 |
| LGC10.1 | 0.6875 | 0.776316 | 0.855769 | 0.857143 | 0.896039604 | 0.701030928 |
| LGC10.2 | 0.875 | 1 | 1 | 1 | 1 | 0.922680412 |
| LGC11.1 | 0.833333 | 0.907895 | 0.99 | 0.945652 | 0.979591837 | 0.779569892 |
| LGC14.1/2 | 0.041667 | 0.763158 | 0.990196 | 0.909091 | 0.942708333 | 0.755102041 |
| LGC14.1/2 | 0.666667 | 0.092105 | 0 | 0.056818 | 0.041666667 | 0 |
| LGC17.1 | 0 | 0.081081 | 0.392157 | 0.378049 | 0.348484848 | 0 |

**Table S7. The most likely combination of parameters of the demographic model from the grid.**

| $N_0$ | $r$ | $K$ | $M$ | $f$ |
|---|---|---|---|---|
| 80 | 0.1 | 2000 | 4 | 2 |

**Table S8. Means and support limits of the demographic model parameters based on interpolation.**

| | mean | 95% support limits |
|---|---|---|
| $\boldsymbol{N_0}$ | 54 | {38, 79} |
| $\boldsymbol{r}$ | 0.12 | {0.09, 0.18} |
| $\boldsymbol{K}$ | 1329 | {940, 1953} |
| $\boldsymbol{K}$ | 3.24 | {2.75, 3.75} |

**Table S9. Proportion of SNPs within each section of the expected range.**

| SNP category | Inside expected range | At or above 0.975 quantile | At or below 0.025 quantile | Above median |
|---|---|---|---|---|
| Control SNPs | 0.92 | 0.04 | 0.04 | 0.53 |
| Spatial outliers | 0.91 | 0.09 | 0.00 | 0.71 |
| Inversions | 0.69 | 0.31 | 0.00 | 1.00 |
